# Flexible Methods for Standard Error Calculation in digital PCR Experiments

**DOI:** 10.1101/2023.09.06.556592

**Authors:** Yao Chen, Ward De Spiegelaere, Wim Trypsteen, David Gleerup, Jo Vandesompele, Olivier Thas

**Affiliations:** Department of Applied Mathematics, Computer Science and Statistics, Ghent University, Ghent, 9000, Belgium; Ghent University Digital PCR Consortium, Ghent University, Ghent, 9000, Belgium; Department of Morphology, Medical Imaging, Orthopaedics, Physiotherapy and Nutrition, Ghent University, Merelbeke, 9820, Belgium; Department of Internal Medicine, Ghent University and University Hospital, Ghent, 9000, Belgium; OncoRNALab, Center for Medical Genetics, Department of Biomolecular Medicine, Ghent University, Ghent, 9000, Belgium; Cancer Research Institute Ghent (CRIG), Ghent, 9000, Belgium; Biogazelle, Ghent, 9052, Belgium; Data Science Institute, I-BioStat, Hasselt University, Diepenbeek, 3590, Belgium; National Institute for Applied Statistics Research Australia (NIASRA), University of Wollongong, Wollongong, 2522, Australia

**Keywords:** digital PCR, standard error, binomial-assumption based methods, non-parametric method

## Abstract

Digital PCR (dPCR) is a highly accurate and precise technique for the quantification of target nucleic acid(s) in a biological sample. This digital quantification relies on the binomial or Poisson distribution to estimate the amount of target molecules based on positive and negative partitions. However, the implementation of these distributions require adherence to underlying assumptions that are often neglected, leading to a suboptimal (too optimistic) variance estimation of the target concentration, especially when considering the multiple sources of variation in experimental dPCR setups. Moreover, these parametric methods cannot be easily used for downstream statistical inference when more advanced analysis are required, such as for copy number variation.

We evaluated the performance of three new statistical methods (BootsVar, NonPVar, BinomVar) in both simulations and real-life datasets for target and variance estimation in dPCR setups while taking into account a combination of commonly observed sources of experimental variability that can interfere with the underlying assumptions of the current parametric methods.

The results demonstrate the capability of the new methods for variance estimation and present a more accurate reflection of the true variability over the classical binomial approach. In addition, these statistical methods are flexible and generic in the way that they work well for the variance estimation of non-linear statistics that work with ratios (e.g. CNV) and for multiplex dPCR setups.

In this study, we provide guidelines when to use the binomial-assumption based methods and when the non-parametric one is better to achieve more accurate variance estimates.

## 1. INTRODUCTION

The use of digital PCR (dPCR) has markedly increased in the last decade. The method, involving massive partitioning of samples, renders each partition an isolated reactor. Few template DNA molecules will end up in the same partition due to the very large number of partitions and sufficiently diluted samples. dPCR has demonstrated many attractive characteristics, for example, inherently high accuracy, no need of calibration standards and providing unsurpassed repeatability [1]. Thanks to this, dPCR is becoming the advised method for absolute DNA or RNA quantification, concentration estimation and copy number variation determination [2, 3, 4].

In a dPCR, the binary outcome, i.e. positive or negative, of a partition is analyzed. Quantification is based on Poisson statistics depending on the counts of positive partitions. Many papers have discussed how to estimate the concentration and the variability of the estimates [5, 6, 7]. Very often a binomial distribution for the number of positive partitions is assumed and a confidence interval is derived accordingly [8]. However, this assumption may not be valid with other sources of variation present [9]. Besides, this method is not very convenient for non-linear functions, e.g. a ratio or a fraction.

The binomial (for singleplex) or multinomial (for multiplex) assumption for the number of positive partitions stands when there is only sampling variation present (approximately Poisson distribution since an individual sample is very small compared to the experiment subject, such as human body, or animals). This is not realistic because other sources of bias and variability will come in before or during the dPCR experiments [9]. For example, there may be pipetting error in mixing the material, or misclassification of partitions after the amplification process. The binomial/multinomial assumption will thus be violated and the existing methods fail to provide correct results.

Variance estimation of a ratio between targets is particularly problematic in dPCR experiments. [10] provided two methods for constructing confidence intervals for CNV. One way is to build a histogram of the ratio by binning and adding up the joint probabilities of all possible combinations of concentrations in each bin of the sampling distributions of the target and reference concentrations. This method has no restriction on the sampling distributions of target or reference concentration, but some prior knowledge on the distribution of the proportions of positive partitions is required. Another method is to use explicit formulas for standard errors and confidence intervals. This method is faster though less accurate than the first one since it assumes that the sampling distribution of the concentration is known. [11] first log transformed the ratio and derived the confidence interval, then the estimates are backtransformed to the original scale. This method makes the assumption that the distribution of the log-ratio is (approximately) normal. The derivation of a formula for the variance estimation of a log ratio can also be analytically difficult. [12] provides a flexible method to account for between-replicate variability by fitting a generalized linear mixed model. This method incorporates the between-replicate variability as a random effect in the statistical model. The replicate effect is described by a normal distribution. So far, this method can be only applied for absolute quantification and CNV.

A limitation of some of the above mentioned methods is that they rely on the assumption of independence of the two variables in the ratio. However, when there is a correlation between the variables, the independent sampling or approximation formulas will be incorrect. For example, dPCR methods aiming to quantify linkage disequilibrium or DNA quality, e.g. the DNA shearing index (DSI), DNA fragmentation [13] will result in a correlation between the measured variables, as these assays are intended to quantify the ratio of linked versus unlinked sequences.

This paper will focus on methods for variance (or standard error) estimation and confidence intervals for the target parameter of a dPCR experiment (e.g. absolute quantifiction, CNV, …). To tackle the above problems, we propose three methods among which two are bootstrap methods that use simulations that mimic the dPCR partitioning process. Two of the methods do not rely on any distributional assumption. Our methods are generic and can be easily applied to many statistics such as ratios and proportions.

## 2. Methodology

### 2.1 A Conditional Bootstrap Method for Standard Errors

The first bootstrap method does not rely on strong distributional assumptions and can be used for standard error calculation when no replicate is available. The method will be referred to as BootsVar. The only assumption that we make, is that the partitioning of molecules is a complete random process. Under these conditions, within a single dPCR run, the conditional distribution of the number of positive partitions, given the total number of partitions *n* and the total number of molecules *m*, is given by Eq.S7 in SI.

Consider the variance estimation of the estimate of the average number of molecules per partition in a single dPCR run. This average is denoted by, λ (which is also the Poisson parameter), and its estimate by 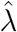 (based on the Poisson approximation, see Eq.S6 in SI).

For a given *m* and *n*, and for a large number *B* (e.g. *B* = 1000), the simulation method in pseudo-code is given as follows:

1. set *l* = 1, and initiate a vector *K*[] of length *B*

2. randomly sample a number from the set {1, 2,. .., *n*} . Each number is sampled with equal chance of 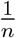 . Repeat the same process *m* times; let **𝒜** denote the set with the unique numbers out of *m* numbers that are sampled

*3. K*[*l*] ← # **𝒜**

4. if *l< B*, then *l* ← *l* +1 and return to step 2; otherwise stop the procedure.

The resulting vector *K*[ ] contains *B* randomly sampled counts of positive partitions from the correct conditional distribution; see Eq.S7 in SI. The proof is straightforward: step 2 randomly samples one partition (out of the *n* partitions) for each molecule (with replacement and hence a partition may contain more than one molecule). The set **𝒜** contains the unique partitions that were sampled and hence its size # **𝒜** (step 3) is the number of positive partitions. Our simulation method basically mimics the partitioning process, for a given *m* and a given *n*.

For each of the *B* elements of the vector *K*[ ], say *K*^*b*^, an estimate of, λ can be computed, say,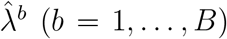. The em irical va ance of the *B* estimates,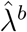 is an approximation of the true variance Var 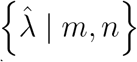. The approximation becomes better as *B* increases. In the limit, as *B* ← ∞ the approximation converges in probability to the true variance (weak law of large numbers).

### 2.2 Binomial Bootstrap for Standard Errors

When *r* replicates are available, we need to account for yet another level of variation: the numbers of molecules loaded on *r* replicated dPCR runs show sampling variability which is caused by sampling (pipetting) from a specimen (a larger volume). The number of molecules is thus a random variable, and will be denoted by *M*_*i*_, for *i* = 1,. .., *r*, with *r* the number of replicates. The distribution function of *M*_*i*_ is denoted by *F*_*M*_ (·; *μ*), with *μ* the expected number of molecules in a sample of constant volume *V*_*d*_. We refer to this random step as *level 1* in Fig. 1.

**Figure 1:**
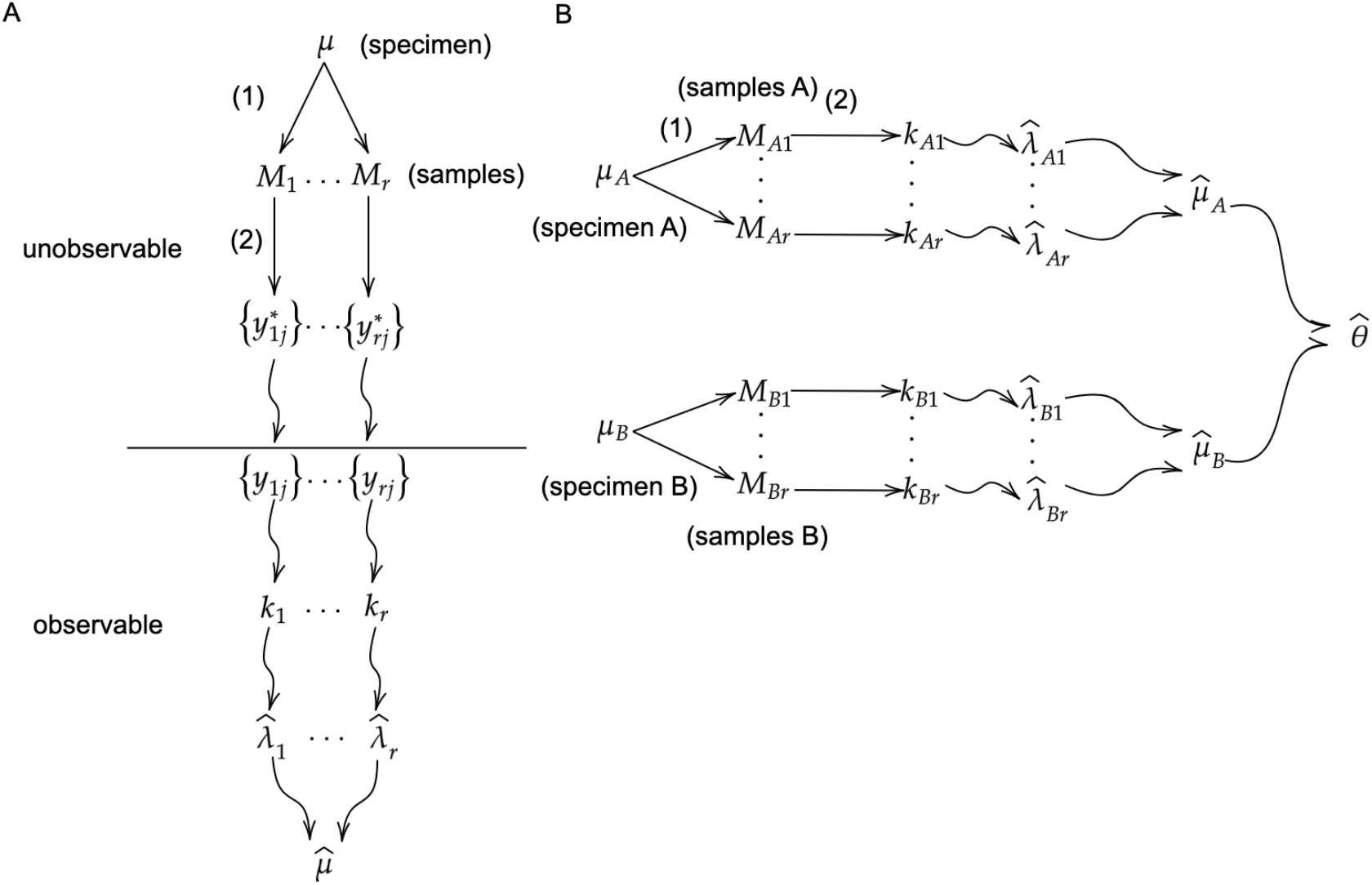
Schematic overview of the random steps (random sampling or partitioning by straight arrows) and calculation steps (label partitions as positive or negative, count the positive partitions and calculate the concentration by curly arrow) in a dPCR experiment with *r* replicates. The numbers 1 and 2 refer to the two random processes (1) random sampling over *r* replicates; (2) random partitioning. (A) single target in detail where *μ* is the concentration of the target (specimen), *M*_*i*_ is the total number of molecules in a sample, 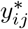 is the exact number of molecules in partition *j* of sample *i, y*_*ij*_ is the binary outcome of a partition (positive or negative), *k*_*i*_ is the count of positive partitions, λ_*i*_ is the average number of molecules in a partition (B) multiple targets where A and B represent different targets

Starting from *M*_*i*_, the molecules are randomly partitioned over *n* partitions, resulting in *K*_*i*_ as the number of positive partitions in replicate *i*. As this is a random step, but conditional on *M*_*i*_, we will write it explicitly as *K*_*i*_ | *M*_*i*_, *n*_*i*_ ∼ *F*_*K*|*Mn*_(·; *M*_*i*_, *n*_*i*_). An important difference with *level 1* is that the distribution *F*_*K*|*Mn*_ is known; it is again Eq.S7 in SI. This random step is referred to as *level 2* in Fig.1. To simplify the notation we will drop the explicit conditioning on the number of partitions *n*_*i*_. The variance of,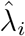 is thus composed of two levels of variability: (1) the random sampling of *M*_*i*_, and (2) given *M*_*i*_, the random partitioning process. Hence, we write Var_*KM*_,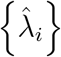 to stress the two random processes.

In this scenario, it is the parameter *μ* that is of interest and needs to be estimated. With *V*_*d*_ the volume loaded in the reaction and *V*_*p*_ the partition volume, this parameter can be estimated as

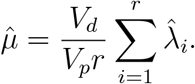

Taking into account both the random partitioning and sampling, the distribution of *K*_*i*_ can be formulated as,

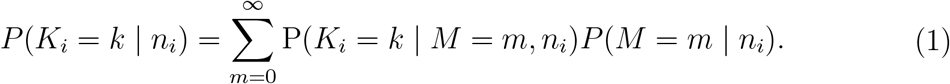

This would be the appropriate distribution for deriving the standard error of,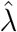, but it can only be used if the distribution of *M* is known.

Upon assuming that the number of molecules *M*_*i*_ are distributed over the replicates as a Poisson distribution with mean *μ*, the distribution of the number of positives *K*, given *n*_*i*_, is approximated by a binomial distribution. This follows from applying Eq.1; see **SI Section Binomial Distribution of Positive Partitions** for a proof. This binomial distribution Binom(*n*_*i*_, π_*i*_) has parameters *n*_*i*_ and 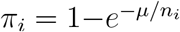, the probability that a partition is positive in replicate *i*.

Since,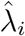 is a (nonlinear) function of *K*_*i*_, the variance of,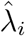(or the variance of a function of the, λ_*i*_, *i* = 1,. .., *r*) can be obtained from this binomial distribution. The conventional method is to apply the delta method, which essentially linearises this nonlinear function (see Eq.S9 in SI). However, here we propose a parametric bootstrap method by resampling numbers of positive partitions from this binomial distribution with the unknown parameter *μ* replaced with its estimate. In particular, this bootstrap algorithm works as follows. With *B* a large number (e.g. *B* = 1000):

1. estimate 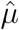 as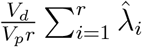, with,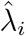 the traditional estimate of, λ_*i*_ based on the Poisson approximation. This equation simplifies to the ordinary mean in most cases.

2. set *i* =1

3. randomly sample *B* observations from Binom(*n*_*i*_, π_*i*_) with 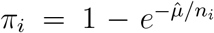 These are denoted as *k*^*b*^, *b* = 1,. .., *B*. For each bootstrap sample *b*, computer 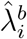

4. based on the *B* estimates,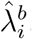, calculate its sample variance 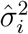

5. if *i*< *r*, then *i* ← *i* +1 and return to step 3; otherwise stop the procedure.

6. average 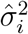 over the *r* replicates

With 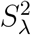 the result of the averaging in step 6, the variance of 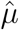 is estimated as 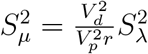

This bootstrap method will be referred to as BinomVar.

Based on the estimated standard error, normal confidence intervals for, λ can be computed [10, 11, 12], which are expected to work well when the estimator,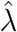 is approximately normally distributed.

### 2.3 A Simple Nonparametric Estimator of the Variance

When one is uncertain about the distribution of the number of molecules *M*, Eq.1 cannot be used.

We propose a simple nonparametric method that does not rely on such a distributional assumption and is therefore expected to be more robust than the previous method (and the delta method) in case of violation of the Poisson assumption (e.g when additional sources of error or variability are involved). Our method only relies on the assumption of random partitioning, i.e. the entry of a molecule to a partition is totally random and the chance is the sa e for ll partitions. In particular, we propose to estimate the variance of 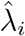 *as*

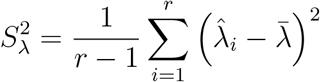

with 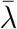 the sample mean of the estimates 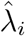 This method will be referred to as NonPVar.

Confidence intervals can be constructed based on the asymptotic normality of the estimator 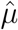, but as a small-sample (i.e. small number of replicates *r*) improvement, we suggest to use quantiles of a *t*-distribution with *r -* 1 degrees of freedom. In particular, a 1 − α confidence interval of *μ* is obtained as

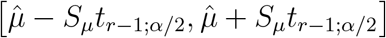

with 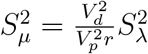.

### 2.4 Additional Sources of Variability and Bias

Both the binomial bootstrap and the delta method assume a binomial distribution for the number of positive partitions *K*, which is derived from a Poisson-distributed *M* . This assumption stands when sampling variability is the sole source of variation in *M* . However, as illustrated in Fig.2, there may be other sources of variability. For example pipetting error, which adds extra variability to the number of molecules loaded on the dPCR, violates the Poisson assumption and therefore invalidating the delta and BinomVar methods.

**Figure 2:**
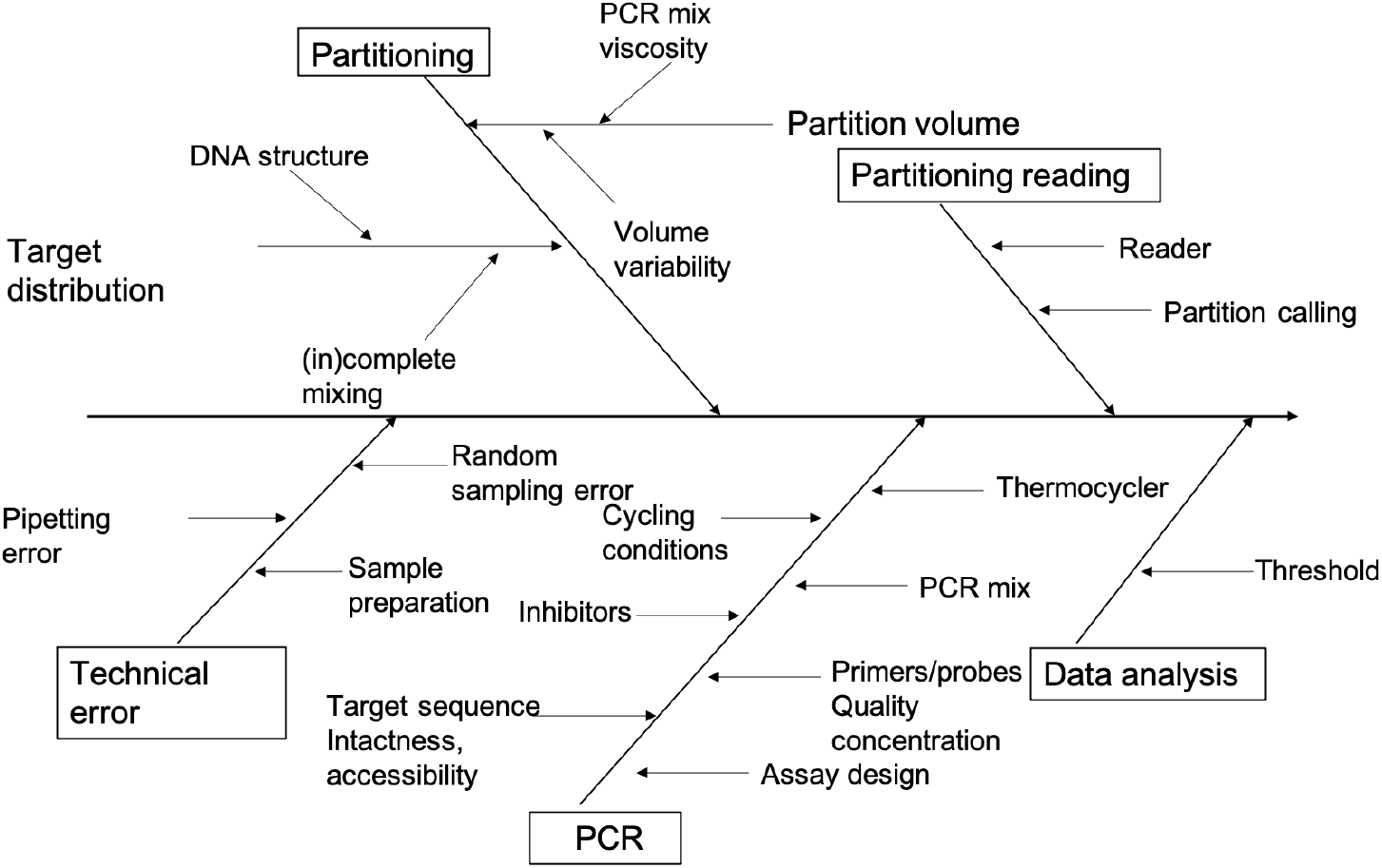
A schematic overview of the dPCR workflow with indications of sources of variability and bias

The NonPVar method, on the other hand, does not make a distributional assumption on *M* . The between-replicate variance is empirically estimated and should therefore cover the sampling variation and other sources of variability. Hence, in situations where the Poisson assumption of *M* is violated, NonPVar is expected to outperform the BinomVar and delta methods.

### 2.5 Absolute Quantification

For absolute quantification, the target parameter is the average number of copies per partition, λ. Here, only one type of target molecules needs to be quantified (see Fig.3A). We also allow for technical replicates.

For all replicates *i* = 1,. .., *r*, we can calculate,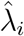 from *K*_*i*_ as in Eq.S6 in SI. Since we have replicates, the final estimator of, λ becomes,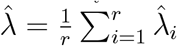, with variance

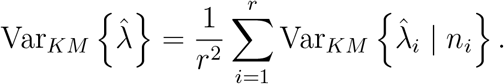

For the estimation of Var_*KM*_,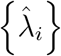 we use the BinomVar procedure as in **Section Binomial Bootstrap for Standard Errors** or nonparametrically estimate the between-replicate variability of,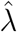 with the NonPVar method as described in **Section A Simple Nonparametric Estimator of the Variance**.

### 2.6 CNV in Singleplex

BinomVar and NonPVar can also be applied to CNV estimation. CNV is defined as large-scale losses and gains of DNA fragments and is one of the major classes of genetic variation [14]. It quantifies how the number of copies of a target gene varies from a reference. In a CNV singleplex set-up, the target (A) and reference (B) molecules are quantified in separate experiments (Fig.3B).

**Figure 3:**
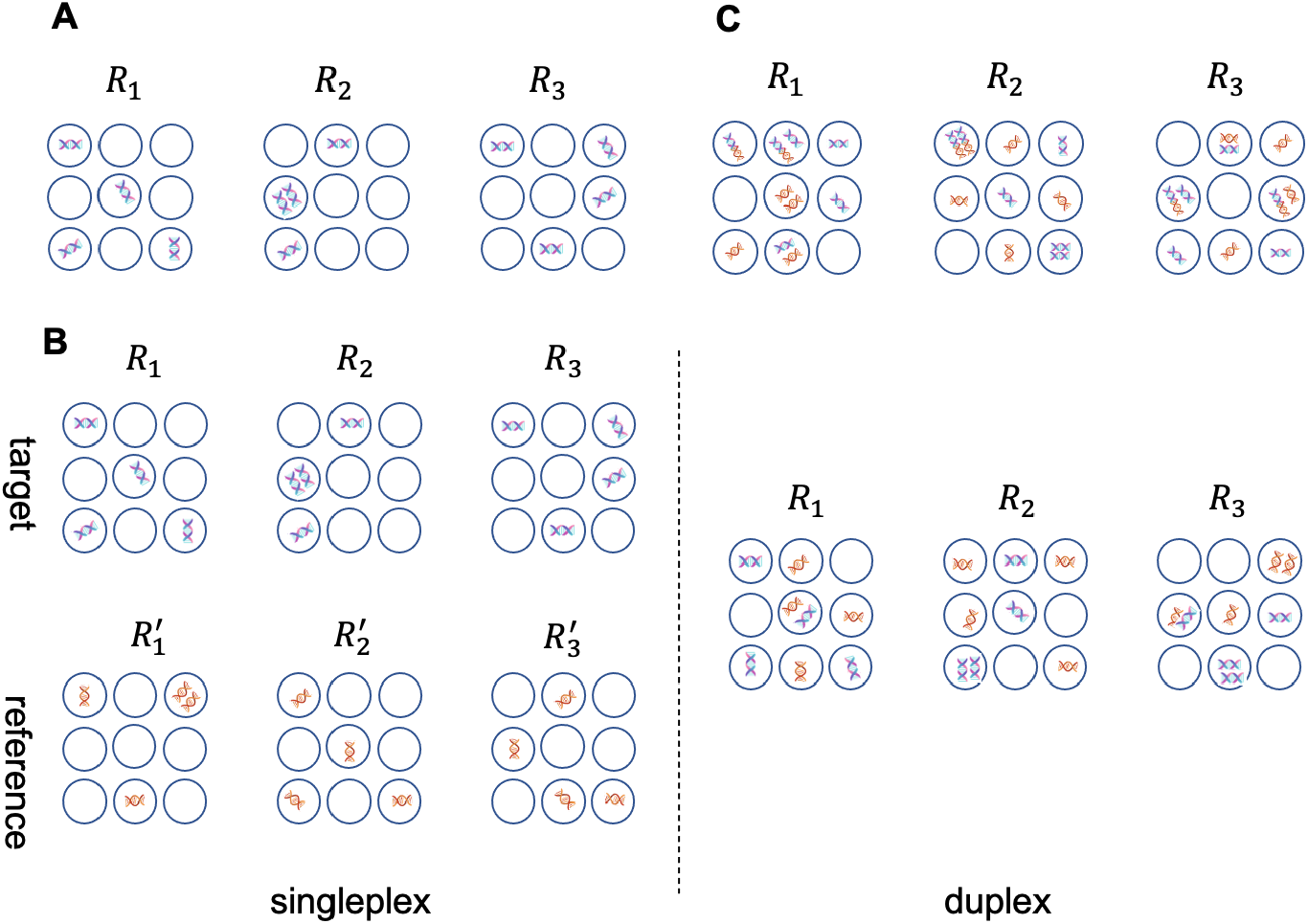
Illustration of the dPCR designs considered in the simulation study: (A) absolute quantification, (B) CNV with singleplex and duplex, and fractional abundance of a mutation in duplex setting, (C) DSI. *R*_1_ up to *R*_3_ refer to replicated experiments

Relying on the same notation as in the previous sections (with obvious extensions), we now consider the estimators

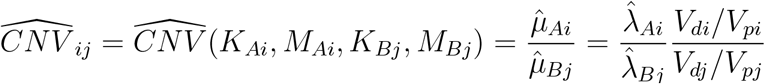

based on replicate *i* (*j*) for molecule A (B). Very often it is reasonable to have *V*_*di*_ equal to *V*_*dj*_ and *V*_*pi*_ equal to *V*_*pj*_. It will turn out to be convenient if we estimate the log-CNV instead,

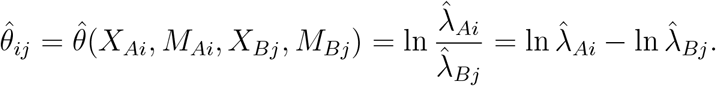

The final estimator of the log-CNV is given by (assuming equal numbers of replicates)

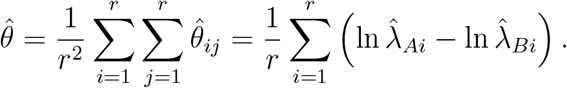

Upon relying on the independence between the singleplex experiments, the variance of 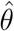 is given by

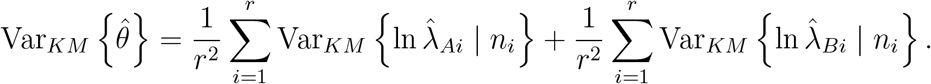

For both terms we apply the same procedures as in **Section Absolute Quantification**. Note that the non-linearity of the log-transformation does not complicate the procedure.

### 2.7 CNV in Duplex, Mutational Load and DSI

In the CNV duplex set-up, the target and reference are typically quantified in the same dPCR run and thus the number of partitions is the same and other sources of errors are shared. There is matching between the replicates for A and B (see Fig.3B). We consider the estimator

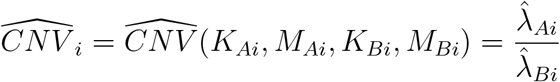

based on replicate *i* for molecules A and B. With the *r* replicates, the final estimator becomes 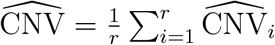 with variance

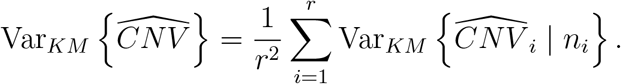

the same procedures as in **Section Absolute Quantification** can also be applied here.

Fractional abundance of a mutation is the component of genetic load attributed to fitness reduction caused by new and recent deleterious mutations [15]. It quantifies the proportion of the mutant type to the total amount of wild and mutant (Fig.3B). The estimator for replicate *i* is as follows,

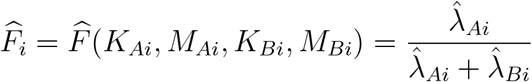

based on replicate *i* for mutant-type A and wild-type B. As it typically is a duplex experiment, both mutant and wild type are quantified in the same experiment. We apply here the same procedures as for the CNV duplex. Note that if the mutant type and wild type are quantified in separate experiments, the estimation procedure will not be the same. NonPVar cannot be applied here due to individual errors and dependency, however, the bootstrap procedure BinomVar is generic and can be extended in this scenario. We can sample from the estimated binomial distributions of the mutant and wild type, calculate the fractional abundance and estimate the variance of the sampled.

DNA shearing index (DSI) is also a proportion and it measures how many DNA fragments are sheared in the preparation stage or during the experiment in order to evaluate gene or genome integrity (Fig.3C). [16] proposed a method for estimating *DSI*, but for such a complicated proportion function, it is difficult to use the delta method or GLMMs [12] to evaluate variability. BinomVar and NonPVar, however, take into account this correlation. For replicate *i*, now consider the estimator,

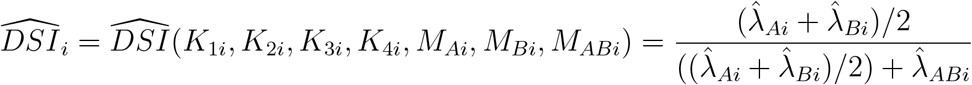

where *K*_1*i*_, *K*_2*i*_, *K*_3*i*_, *K*_4*i*_ are the partition numbers for single positive A, double positive, double negative and single positive B of replicate *i* respectively, and,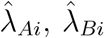 and,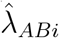 are the corresponding estimated average numbers of target molecules per partition. The same procedures as for CNV in duplex can be applied here.

### 2.8 Simulation Study

The new methods are evaluated and compared to a few competitor methods in a simulation study. Realistic simulations of numbers of positive partitions allow us to assess methods under various scenarios. Moreover, by repeatedly simulating data for a single scenario, the statistical properties of the methods can be evaluated.

We start with a setup of a single replicate to investigate the estimator of the conditional variance given a fixed *m* and *n*, as described in **Section A Conditional Bootstrap Method for Standard Errors**. For multiple replicates, we follow the simulation pipeline of [9]; see also Fig. 2. In the first scenario the number of molecules *M* is randomly sampled from a Poisson distribution, and next, given *M*, the number of positive partitions are generated by random partitioning of the molecules over *n* partitions. Subsequent scenarios add additional sources of variation and bias to the data generating process, as illustrated in Fig. 2. See **SI Section Simulation Settings** for a detailed description of the scenarios.

These scenarios were used for evaluating the performance of the methods for the following target parameters: absolute quantification and CNV, both in singleplex, and for CNV, fractional abundance of a mutation and DSI, all in duplex. As competitor methods we included the delta and the GLMM [12] methods for absolute quantification and CNV in singleplex and duplex. For the fractional abundance of a mutation and DSI, however, there are no competitor methods to our knowledge.

For each scenario, the methods were evaluated based on 1000 simulation runs, and within each run the bootstrap methods were applied with 1000 bootstrap runs. The performance of the variance estimators is evaluated in terms of the bias (relative bias or absolute bias w.r.t. the true variance), and the performance of the 95% confidence intervals is assessed in terms of the empirical coverage (i.e. the relative frequency, over the 1000 simulation runs, that the true target parameter falls within the CI).

### 2.9 Implementation

The data analysis was performed in R (version 4.2.0). The codes can be found at https://github.com/emmachenlingo/dpcr-flexible-methods-for-standard-error-calculatiotree/digital-PCR. A Shiny web application is also available at https://dpcr-ugent.shinyapps.io/variance_estimate/ that does not require knowledge about programming, and relies on a spreadsheet-like data input and point and click interface.

## 3. RESULTS and DISCUSSION

### 3.1 Without Replicate

Through the simulation runs, the total number of molecules (*m*) and partitions (*n*) are held constant.

The results (see Table S1 in SI) show that, except for very low concentrations (≈ 0.005 molecules per partition on average), the BootsVar variance estimator is nearly unbiased. The BinomVar, delta and GLMM methods overestimate the variance estimates because these methods are all based on the binomial assumption of the positive partitions, which does not stand with a fixed *m*. Consequently, BinomVar, Delta and GLMM give too wide confidence interval (see Fig.S1 in SI). We also compared the computation time of those methods (see Table S2 in SI). BootsVar, with embedded 1000 Bootstrap iterations, will take longer time than other methods. But for small-to-medium size dataset, it can still handle.

### 3.2 With Replicates

For absolute quantification, all methods give nearly unbiased estimators of, λ, except in the presence of partition size variation and misclassification (see Fig.S2 to S7 in SI). The effect of partition size variation is limited, but misclassification has a larger impact; this agrees with the findings of [9]. Regarding the latter, assuming constant false positive and false negative rates for all replicated experiments, the estimates will be consistently biased downwards or upwards depending on the concentration.

Fig.4A shows that with only sampling variability and random partitioning, all methods give good CIs. In the presence of pipetting error, both the delta method and BinomVar cover the true value with a probability of less than 50% when, λ *>* 0.5 (Fig.4B). The NonPVar method is more robust against such errors and performs the best. The GLMM method is the runner-up.

**Figure 4:**
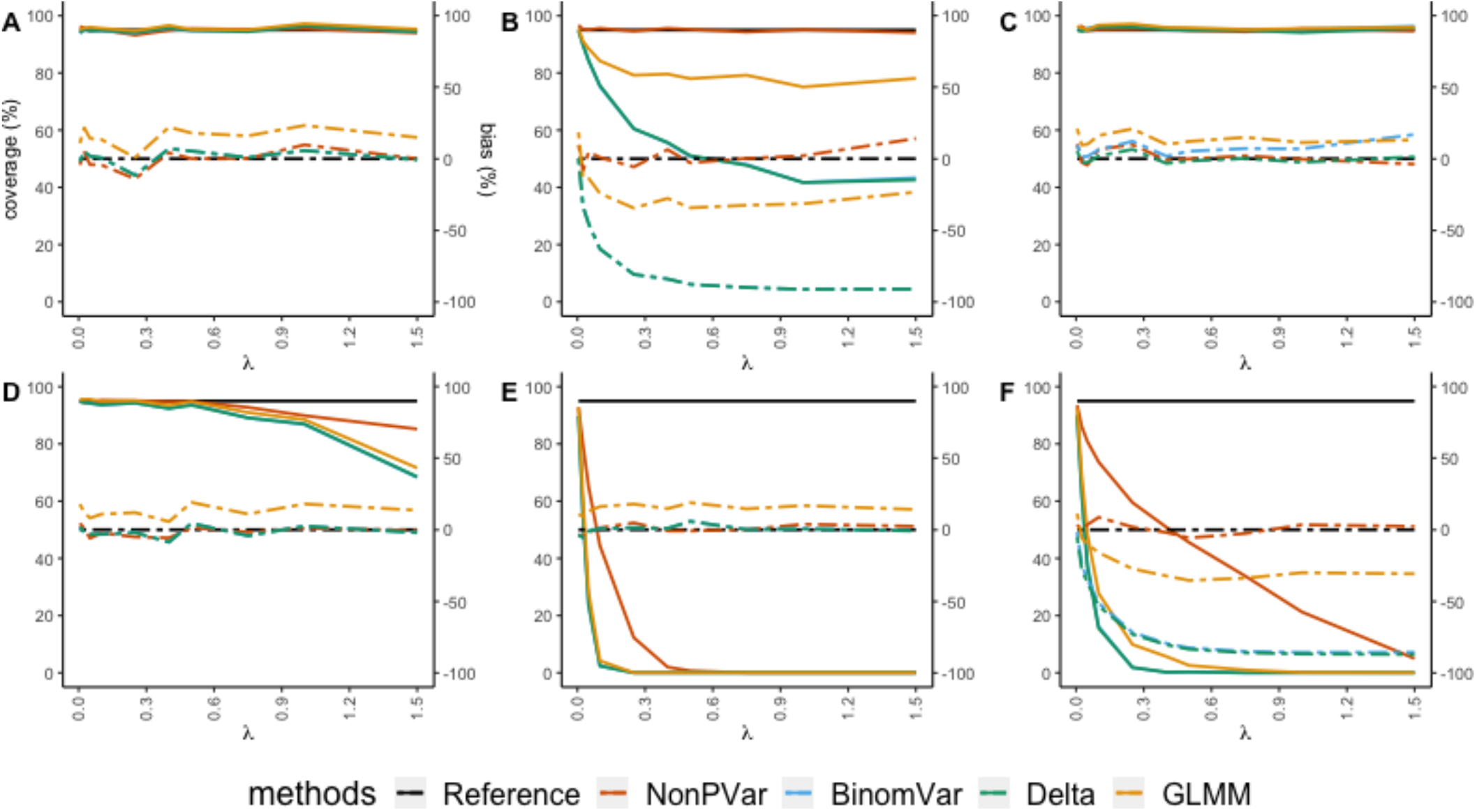
Empirical coverage of the 95% CIs (solid lines, left axis) and relative bias (dashed lines, right axis) for absolute quantification in different scenarios. X-axis represents varying concentration of target molecules from low to high. (A) only sampling variation and random partitioning with 3 replicates (B) 3% pipetting error (C) 20% partition loss (D) coefficient of variation of 10% in partition size (E) misclassification with 0.01% false positive rate and 5% false negative rate (F) all variation included. The reference (black solid line) for empirical coverage is set at 95%. The constructed CIs are supposed to cover the true values in 95% of the time. The closer other solid lines are to this reference, the better the CIs are. The reference (black dashed line) is set at 0%. The closer other dashed lines are to this reference, the lower the relative bias is.

In terms of the variance estimation, NonPVar has very low relative bias in all scenarios. With additional pipetting error, the NonPVar variance estimator is much less biased than its competitors. The absolute bias and the distribution of the variance estimates (over the simulation runs) are also checked (see Fig.S8 to S19 in SI). However, the boxplots of the variance estimates show that the NonPVar estimates have a larger variance than the alternative methods. This is probably because NonPVar estimates the variance empirically based on only a limited number of replicates, whereas the other methods rely on a distributional assumption for the variance estimation. With additional pipetting error, only variance estimates by NonPVar and GLMM are close to the true value. The BinomVar and Delta methods underestimate the variance. Partition size variation and misclassification have an impact on the coverage of confidence interval, especially misclassification. All methods fail to cover the true value when partitions are misclassified and the error is consistent through all replicates.

For CNV in singleplex, target and reference molecules are quantified separately. That means that additional sources of variability such as pipetting error will be different for target and reference genes. Fig.S20 in SI shows similar pattern as in absolute quantification. The results suggest that NonPVar performs at least as good as the other methods in terms of empirical coverage, while its relative bias remains quite low.

In multiplexing experiment, all target DNA molecules are quantified within the same reaction, and thus additional sources of variation apply equally to them.

In the simulations for DSI, the concentration and intactness percentage are varied from low to high. Results in Fig.5 (see also Fig.S61 and S62 in SI) show that the effect of pipetting error, which commonly have a big impact on the variation and confidence interval of the estimates in absolute quantification or CNV singleplex setup, is now canceled out. The empirical coverage of BinomVar is very high, even with the presence of pipetting error. The relative biases of NonPVar and BinomVar are both close to 0. However, the NonPVar estimates are less precise (see Fig.S49 to S60 in SI) because of its empirical nature for estimating the variance (in the absence of distributional assumptions).

**Figure 5:**
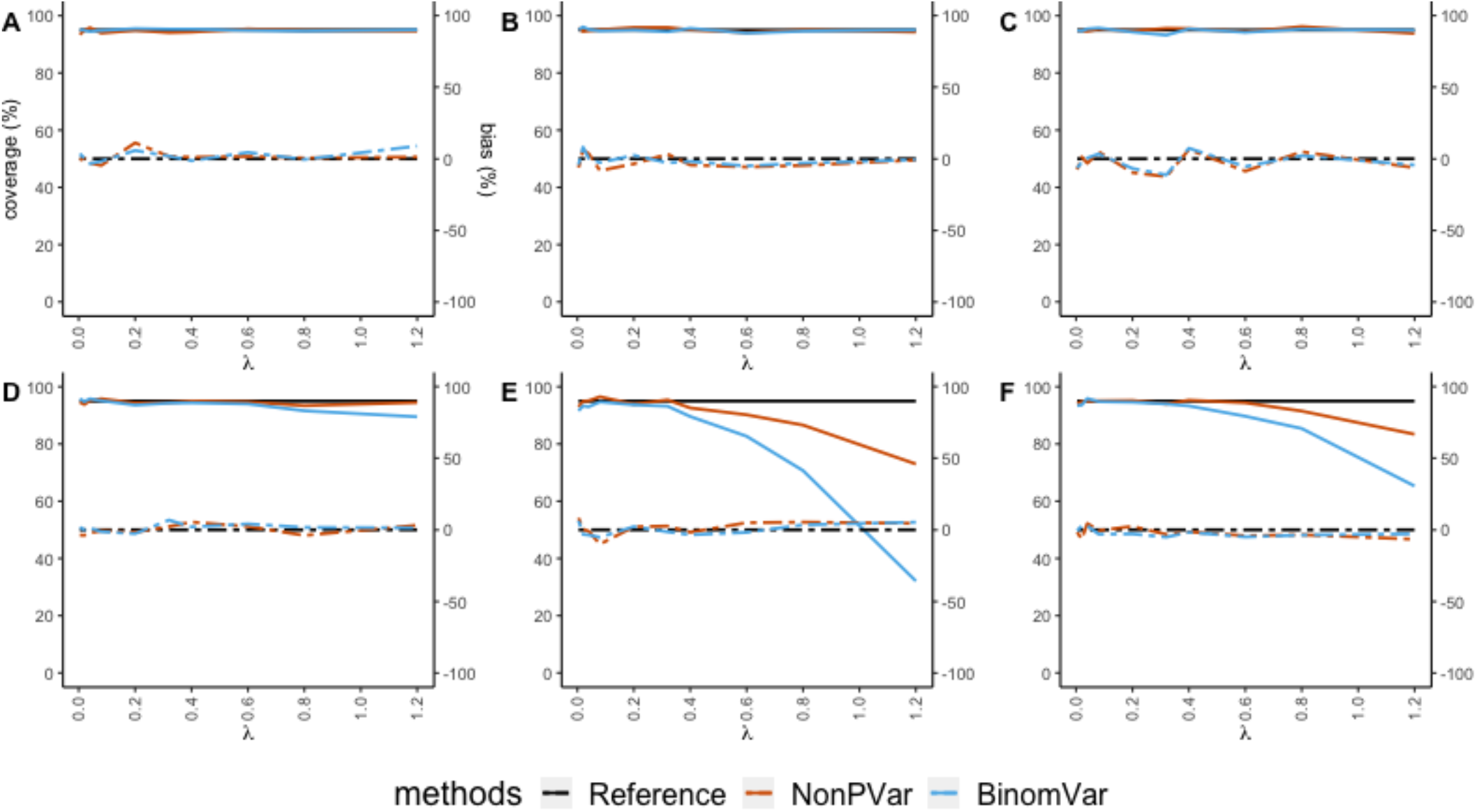
Empirical coverage of the 95% CIs (solid lines, left axis) and relative bias (dashed lines, right axis) for low DSI (=20%, that is, 20% of the target molecules got sheared) in different scenarios. X-axis represents varying concentration of intact molecules from low to high. (A) only sampling variation and random partitioning with 3 replicates (B) 3% pipetting error (C) 20% partition loss (D) coefficient of variation of 10% in partition size (E) misclassification with 0.01% false positive rate and 5% false negative rate (F) all variation included. The reference (black solid line) for empirical coverage is set at 95%. The constructed CIs are supposed to cover the true values in 95% of the time. The closer other solid lines are to this reference, the better the CIs are. The reference (black dashed line) is set at 0%. The closer other dashed lines are to this reference, the lower the relative bias is.

For fractional abundance of a mutation, results in Fig.S40 in SI show that without classification error, the performance of NonPVar and BinomVar is quite comparable. With misclassified partitions, the empirical coverage of BinomVar is similar to that of NonPVar in low or medium concentration scenarios, but it is considerably lower in the high concentration scenarios.

Results for CNV duplex (see Fig.S39 in SI) show similar patterns as DSI and fractional abundance of a mutation. The effect of misclassification is not found, because in this scenario the numbers of target and reference genes coincide and so the effect is simply canceled out.

### 3.3 Case Study

In the case study, we investigated two types of real-life data: CNV in singleplex and mutations. To assess the variance, replicates of samples were analyzed.

The Dataset of mutations comes from [17] and 3 types of samples were included: (i) patient samples with a very high mutation load (samples 1– 13); (ii) homoplasmic wild-type samples from a healthy volunteer (samples 14–16); and (iii) samples undergoing nuclear transfer, thus carrying a low mutation load due to mtDNA carry-over (samples 17–23). 3 technical replicates were obtained for 6 samples.

Results in Fig.6 show that for some samples, the two confidence intervals are quite different and for others, they are comparable. In a sense of absolute value, the difference is not very large. The sample concentrations of this dataset are low. And from the simulation results, it is observed that in low concentration, the random sampling variability is dominating and other sources of error, such as misclassification whose effect will not cancel out, do not have a big impact yet. In this scenario, BinomVar would give more precise variance estimates, thus should be preferred.

**Figure 6:**
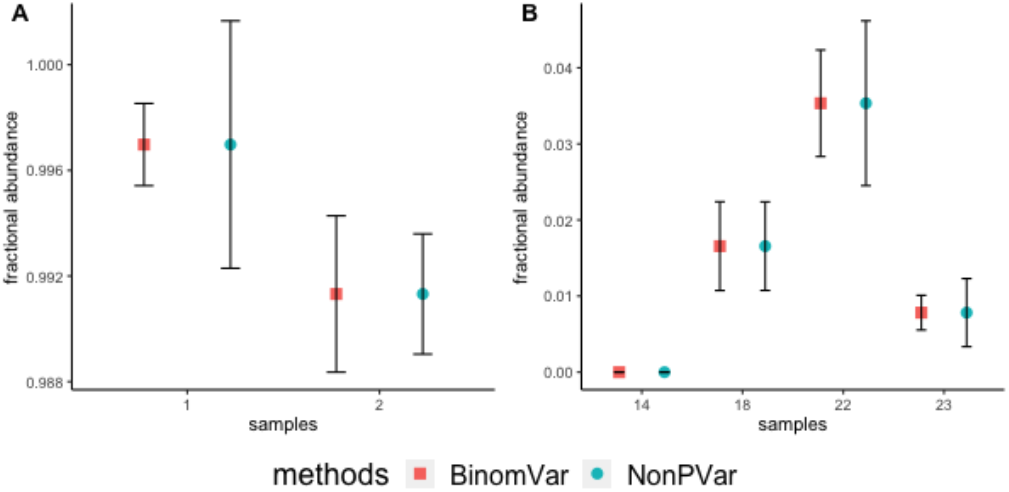
Fractional abundance of a mutation in sample 1,2,6,14,18 and 23 from the dataset in [17] estimated with BinomVar and NonPVar.

For the CNV dataset, overall, confidence intervals given by BinomVar, delta method and GLMM are close (see Fig.S63 and S64 for more details). One possibility is that those three methods behave more similarly in lower range of concentrations (the average number of target molecules per partition in sample 15 ranges from 0.097 to 0.19 over the genes with a mean of 0.13). In the low concentrations, the number of molecules in a sample is likely still to follow a Poisson distribution, so BinomVar, Delta and GLMM would be recommended. The running time (Table S3 in **Section Real-life data analysis**) shows that all methods except for GLMM are really fast. However, for small-to-medium size datasets, computation time of GLMM is quite acceptable.

### 3.4 Impact of Different Sources of Variability

The types of dPCR data are generated in different ways. For singleplex experiments, there is no dependency among the different types of molecules. For duplex or higher order multiplexing, there is a natural dependency. No dependency implies that we should consider variability such as pipetting error separately while dependency means those sources of errors are shared for all target molecules. Both the NonPVar and BinomVar methods are generic and manage to give a good estimate of the confidence interval with or without dependency.

In the simulation study we show that the binomial assumption of positive partitions is violated with the presence of pipetting error. Confidence intervals relying on this assumption therefore may fail to cover the true value most of the time. The NonPVar method is more robust against such error. Note that different sources of variability have different impacts on the variance estimates (Fig.4, 5, S20, S39, S40). In absolute quantification and the CNV singleplex set-up, ignoring pipetting error will result in underestimation of the variance and inaccurate confidence interval. For CNV duplex, fractional abundance of a mutation or DNA shearing index, most of the additional errors cancel. The effect of varying partition size does not disappear even in duplex or multiplex set-up, because all methods make use of Poisson statistics to calculate, λ which is built upon the assumption of random partitioning and equal chance to enter any partition. This assumption is violated if the partition volume is not constant. Our finding aligns with previous studies [18, 19, 20, 9, 8, 21]. It indicates that information about partition-volume is needed to reduce the estimation bias. We also notice that the impact of misclassification cancels out only when the numbers of target and reference molecules has a ratio of 1:1 or when half of the intact fragments get sheared. When the ratio is not close to 1, the effect of misclassification is no longer negligible. This indicates a clear need to use a good partition classification method for higher multiplexing.

### 3.5 Suggestion on Variance Estimation Methods

NonPVar is a data-driven method and does not rely on a distributional assumption of positive partitions. Errors are inferred from the data. Since the method makes use of replicates, the estimation accuracy depends on the number of replicates. Particularly the NonPVar method, despite being unbiased on many scenarios, requires a sufficient number of replicates to give precise variance estimates. The sample size can be calculated based on the required precision [22]. The BinomVar method gives more precise estimates, because it relies on the Poisson assumption for the sampling distribution of the number of molecules over the replicates. The price that BinomVar pays is that it becomes less robust when this distributional assumption is violated.

Results also show that in low concentration scenarios, other sources of errors such as pipetting error do not have a very big impact and BinomVar is the better choice. In summary, we would suggest choosing estimation methods by the type of experiments, concentration levels and replicates number, see Table 1.

**Table 1:** Suggestion on estimation methods. The choice of methods also depends on the sample size. If the precision requirement is met, NonPVar will be as good as other methods in duplex/multiplex scenarios. Note there is no exact threshold to define low or high concentration levels. According to our simulation studies, λ*<* 0.1 can be considered as low concentration level. However, it also depends on the quality of the data. If there is less pipetting error and targets are accurately quantified, then the threshold should be higher.

| Concentration levels | Types of experiment | Recommended methods |
| --- | --- | --- |
| **Low** | Singleplex | BinomVar, GLMM or Delta method |
|  | Duplex/Multiplex | BinomVar, GLMM or Delta method |
| **High** | Singleplex | NonPVar |
|  | Duplex/Multiplex | Depends on the classification. If the clusters are well-separated, then BinomVar, GLMM or Delta method. Otherwise, misclassification error will be high and NonPVar is a better choice. |

## 4 CONCLUSION

We propose three new methods (BootsVar, NonPVar and BinomVar) for variance estimation with dPCR data. BootsVar mimics the random partitioning process in the dPCR experiment through a bootstrap procedure and can be used for variance estimation when replicates are not available. The NonPVar method accounts for inner variability (induced by the technology itself) and between-replicate variability (sampling variability and also other sources of variation). BinomVar uses a Bootstrap procedure that is based on a binomial assumption. The methods circumvent the mathematical formulation of the variance formula, which makes them generic and easy to use. The NonPVar method often outperforms the other methods in terms of empirical coverage of the confidence interval and the relative bias. However, it gives less precise estimates than the BinomVar method. We suggest that in the lower range of concentrations, the distributional assumptions based methods (BinomVar, the delta method or GLMM) are used, and for the high concentrations or in the presence of other sources of variability, the NonPVar method should be used.

## Acknowledgement

This work was funded by the Ghent University Special Research Fund, BOF (grantnr: 01IO0420).

## Supporting Information

The Supporting Information is available in SI file.

