## Supplementary material for "Flexible Methods for Standard Error Calculation in digital PCR Experiments": SI

**2 Simulation Settings** **11**

**3 Conditional Variance** **12**

**4 Absolute Quantification** **13**

**5 Copy Number Variation** **23**

**8 Real-life Data Analysis** **47**

### 1 Methodology

#### 1.1 Single Replicate

##### 1.1.1 Probabilistic Framework

A typical dPCR data analysis starts from the end-point fluorescence after a fixed number of amplification cycles. The raw continuous fluorescence levels are transformed to binary (digital) observations after applying a threshold. In particular, when the end-point fluorescence exceeds the threshold, the partition is labelled positive, otherwise negative. Let $n$ denote the number of partitions. The relation between the binary outcome $Y_j$ of partition $j$ and the unobserved count of the target molecule $Y_j^*$ in that partition can be formulated as ($j = 1, \ldots, n$)

$$Y_j = \min\left(Y_j^*, 1\right) = \begin{cases} 0 \text{ if } Y_j^* = 0 \\ 1 \text{ otherwise} \end{cases}, \tag{S1}$$

i.e. $Y_j$ is 0 if there are no copies and it is 1 if there is at least one copy.

As the total number of molecules ($m$) in the sample is fixed and the entry of a molecule into a partition is random, the counts $Y_j^*$ follow a binomial distribution with distribution function

$$\mathrm{P}(Y_j^* = y \mid m, n) = \binom{m}{y} \left(\frac{1}{n}\right)^y \left(1 - \frac{1}{n}\right)^{m-y}. \tag{S2}$$

When $n$ is large enough, this binomial distribution can be approximated by a Poisson distribution with parameter $\lambda = m/n$ which can be interpreted as the average number of target molecules per partition. The distribution function of this Poisson distribution is given by

$$\mathrm{P}(Y_j^* = y \mid \lambda) = \frac{\lambda^y e^{-\lambda}}{y!}. \tag{S3}$$

The $\lambda$ parameter can be directly estimated from the digital outcomes, because $Y_j^* = 0$ if and only if $Y_j = 0$ (i.e. a partition containing no molecules is a negative partition). Upon using this relationship the Poisson distribution gives

$$\mathrm{P}\left\{Y_j^* = 0 \mid \lambda\right\} = \frac{\lambda^0}{0!}\exp(-\lambda) = \exp(-\lambda). \tag{S4}$$

Hence the relationship

$$\lambda = -\log \mathrm{P}\left\{Y_j^* = 0 \mid \lambda\right\} = -\log \mathrm{P}\left\{Y_j = 0 \mid \lambda\right\}. \tag{S5}$$

Since the digital outcomes $Y_j$ are observed, the probability of a negative partition, $\mathrm{P}\{Y_j = 0 \mid m, n\}$, can be estimated by $1 - K/n$, where $K = \sum_{j=1}^{n} I(Y_j = 1)$ is the number of positive partitions. The estimate of $\lambda$ thus becomes

$$\hat{\lambda} = -\log\left(1 - \frac{K}{n}\right). \tag{S6}$$

This parameter estimate is crucial in most of the dPCR applications. For example, in absolute quantification the concentration of the target is estimated as $\hat{\lambda}/V_p$, with $V_p$ the (average) volume of a partition. Another example: CNV is based on the ratio of two $\hat{\lambda}$'s; e.g. one for the target and one for the reference.

The imprecision of the estimate $\hat{\lambda}$ can be expressed as its standard error $\mathrm{se}(\hat{\lambda} \mid m, n)$ or its variance $\mathrm{Var}\left\{\hat{\lambda} \mid m, n\right\} = \mathrm{se}^2(\hat{\lambda} \mid m, n)$. The focus of this paper is on the estimation of this variance. Imprecision can also be expressed as a confidence interval (CI) of $\lambda$. If the sampling distribution of $\hat{\lambda}$ is approximately normal, then an approximate 95% CI can be calculated as $\hat{\lambda} \pm 1.96 \times \widehat{\mathrm{se}}$, with $\widehat{\mathrm{se}}$ the estimated standard error. Later we will also propose another method for CI calculations. Since $\hat{\lambda}$ is a function of the number of positives $K$, the variance of $\hat{\lambda}$ depends on the distribution of $K$.

Suppose the number of molecules in the sample ($m$) is fixed, as well as the total number of partitions in the dPCR run ($n$). Then the probability of $k$ partitions being positive is given by

$$P(K = k \mid m, n) = \frac{n! S_2(m, k)}{(n-k)! n^m}, \tag{S7}$$

where $S_2(m, k)$ refers to the Stirling number of the second kind, which is the number of ways to partition $m$ objects into $k$ non-empty subsets (Graham et al., 1989). Since $m$ and $n$ are considered fixed, this distribution reflects the variability in the number of molecules per partition as a consequence of the random partitioning. Although with this distribution function and with basic probability calculus it is possible to find the variance of $\hat{\lambda}$, these calculations are hard. Instead the distribution of $K$ is assumed approximately a binomial distribution:

$$K \mid n, \lambda \sim \text{Binom}(n, 1 - \exp(-\lambda))) \tag{S8}$$

in which $1 - \exp(-\lambda)$ is the probability of a positive partition (see Equation (S4)). Based on the binomial we have $\text{Var}\{K \mid \lambda, n\} = n \exp(-\lambda)(1 - \exp(-\lambda))$ and hence the conditional variance of $\hat{\lambda}$, which is the nonlinear function (Eq.S6) of $K$, can be approximated upon using the delta method (based on a linearisation of the nonlinear log-function).

##### 1.1.2 Delta Method for the Standard Error

The delta method is a well established method in statistics for approximating the variance of estimators that are nonlinear functions of the sample observations. It is based on a first order Taylor expansion of this nonlinear function.

This method has been applied to the estimator $\hat{\lambda}$ of Eq.S6, resulting in

$$\text{Var}\left\{\hat{\lambda} \mid n, \lambda\right\} \approx \frac{\pi}{n(1-\pi)} \tag{S9}$$

with $\pi = 1 - \exp(-\lambda)$ the probability of a positive partition. With $\hat{\pi} = 1 - \exp(-\hat{\lambda}) = K/n$ an estimator of $\pi$, the conditional variance of $\hat{\lambda}$ can be computed by substituting $\pi$ with $\hat{\pi}$ in Eq.S9, resulting in the variance estimator $K/[n(n-K)]$.

Based on the delta method, Whale et al. (2012) gives an approximation of the variance of CNV estimators based on two independent singleplex experiments. The numbers of positive partitions of both the target and reference are considered to follow the binomial distribution (Eq.S8). The delta method is applied to the log transformed CNV estimate, resulting in the approximation

$$\text{Var}\left\{\log \frac{\hat{\lambda}_t}{\hat{\lambda}_r} \mid n, \lambda_t, \lambda_r\right\} \approx \frac{(1-\exp(-\lambda_t))}{(n \times \lambda_t^2 \exp(-\lambda_t))} + \frac{(1-\exp(-\lambda_r))}{(n \times \lambda_r^2 \exp(-\lambda_r))}, \tag{S10}$$

where $\lambda_t$ and $\lambda_r$ refer to the target and reference, respectively. An estimator of this variance is obtained by replacing the $\lambda$ parameters by their estimators (Eq.S6). A disadvantage of this approach is that it only gives a variance estimate of the log-CNV; it cannot be accurately backtransformed to the original CNV scale. If the estimate is used for the calculation of a CI of the log-CNV, then the boundaries of this interval can be correctly backtransformed to the boundaries of a CI of the CNV by exponentiating these bounds.

##### 1.1.3 Distribution of positive partitions

Given a fixed number of molecules $m$, the number of positive partitions $k$ can be modeled by the following distribution,

$$\begin{aligned} & P\left(K=k \mid m, n\right) \\ = & \frac{n! S_2(m,k)}{(n-k)! n^m} \\ = & \frac{n! \left\{ {m \atop k} \right\}}{(n-k)! n^m} \\ = & \frac{n!}{(n-k)! n^m} \frac{1}{k!} \sum_{i=0}^{k} (-1)^i \left( \begin{array}{c} k \\ i \end{array} \right) (k-i)^m \end{aligned} \tag{S11}$$

where $S_2(m,k)$ refers to Stirling number of the second kind, which is the number of ways to partition $m$ objects into $k$ non-empty subsets. This equation is also mentioned in Debski and Garstecki (2016). For the sake of computation, $S_2(m,k)$ is approximated as,

$$\left\{ \begin{array}{c} m \\ k \end{array} \right\} \sim \sqrt{\frac{v-1}{v(1-G)}} \left( \frac{v-1}{v-G} \right)^{m-k} \frac{k^m}{m^k} e^{k(1-G)} \left( \begin{array}{c} m \\ k \end{array} \right) \tag{S12}$$

where $G = -W_0\left(-ve^{-v}\right), v = m/k$, and $W_0(z)$ is the main branch of the Lambert W function (Temme, 1993). Eq.S11 is also log-transformed to deal with factorial and power function.

$$\begin{aligned} l = & \sum_{i=1}^{n} log(i) - \sum_{i=1}^{n-k} log(i) - mlog(n) + \frac{1}{2} log(\frac{v-1}{v(1-G)}) + (m-k) log(\frac{v-1}{v-G}) \\ & + mlog(k) - klog(m) + k(1-G) + \sum_{i=1}^{m} log(i) - \sum_{i=1}^{m-k} log(i) - \sum_{i=1}^{k} log(i) \end{aligned} \tag{S13}$$

this equation requires integer $n, m$ and $k$.

##### 1.1.4 Binomial Distribution of Positive Partitions

Let $M_{pj}$, i.i.d Poisson($\lambda$), denote the number of molecules in partition $j$.

The number of molecules in the sample, $M$, can then be written as

$$M = \sum_{j=1}^{n} M_{pj}.$$

Since all $M_{pj}$ are i.i.d. Poisson distributed, we find

$$M \sim \text{Poisson}(n\lambda).$$

Note that these Poisson distributions are marginal distribution in the sense that we did not condition on the number of molecules.

Let $K_{pj}$ denote the 0/1 indicator for partition $j$ to be positive. Then the number of positive partitions $K = \sum_{j=1}^{n} K_{pj}$. These $K_{pj}$ can be described by a Bernoulli distribution with parameter $\pi = P(K_{pj} = 1)$, which is the probability that a partition is positive. Since the hit of a partition by a molecule is completely at random, all $K_{pj}$ are identically distributed. Thus, $K_{pj}$ i.i.d. $B(\pi)$, and

$$K = \sum_{j=1}^{n} K_{pj} \sim \text{Binom}(n, \pi).$$

The distribution of $K$ can be deconvoluted as,

$$\text{P}(K = k) = \sum_{m=0}^{\infty} \text{P}(K = k \mid M = m) g(m).$$

We find

$$\begin{aligned}
\text{P}(K = k) &= \sum_{m=0}^{\infty} \frac{n!}{(n-k)! n^m} mk \frac{(n\lambda)^m}{m!} e^{-n\lambda} \\
&= \frac{n!}{(n-k)!} e^{-n\lambda} \sum_{m=0}^{\infty} mk \frac{\lambda^m}{m!} \\
&= \frac{n!}{(n-k)!} e^{-n\lambda} \frac{1}{k!} \left(e^{\lambda} - 1\right)^k \\
&= \frac{n!}{(n-k)! k!} \left(e^{-\lambda}\right)^{n-k} \left(1 - e^{-\lambda}\right)^k .
\end{aligned}$$

We have used the *exponential generating function*,

$$\lim_{u \to \infty} \sum_{m=0}^{u} mk \frac{\lambda^m}{m!} = \frac{1}{k!} \left(e^{\lambda} - 1\right)^k .$$

The final expression for $P(K = k)$ shows the binomial distribution function $\text{Binom}(n, \pi)$ with $\pi = 1 - e^{-\lambda}$ the probability that a partition is positive. Or, equivalently,

$$\lambda = -\log(1 - \pi)$$

with $1 - \pi$ the probability that a partition is negative. This is exactly the equality that is used in the classical Poisson approach.

In conclusion, the distribution of positive partitions is compatible with the Poisson and binomial distributions which arise from the classical approach.

##### 1.1.5 Expected value of positives

We know that

$$\begin{aligned} \left\{ \begin{array}{c} m+1 \\ k \end{array} \right\} &= k \left\{ \begin{array}{c} m \\ k \end{array} \right\} + \left\{ \begin{array}{c} m \\ k-1 \end{array} \right\} \\ \sum_{k=1}^{m} \left\{ \begin{array}{c} m \\ k \end{array} \right\} n^{\underline{k}} &= n^m \end{aligned} \tag{S14}$$

where $\left\{ \begin{array}{c} m+1 \\ k \end{array} \right\}$ stands for $S_2(m+1, k)$, and $n^{\underline{k}}$ is the falling factorial, $n(n-1)...(n-k+1)$. Then

$$
\begin{aligned}
&E\left(k \mid m\right)\\
=&\sum_{k=1}^{m} k\frac{n!S_2(m,k)}{(n-k)!n^m}\\
=&\frac{1}{n^m}\sum_{k=1}^{m} k \left\{ \begin{matrix} m \\ k \end{matrix} \right\} n^{\underline{k}}\\
=&\frac{1}{n^m}\sum_{k=1}^{m} \left( \left\{ \begin{matrix} m+1 \\ k \end{matrix} \right\} - \left\{ \begin{matrix} m \\ k-1 \end{matrix} \right\} \right) n^{\underline{k}}\\
=&\frac{1}{n^m}\left( \sum_{k=1}^{m} \left\{ \begin{matrix} m+1 \\ k \end{matrix} \right\} n^{\underline{k}} - \sum_{k=1}^{m} \left\{ \begin{matrix} m \\ k-1 \end{matrix} \right\} n^{\underline{k}} \right)\\
=&\frac{1}{n^m}\left( \sum_{k=1}^{m+1} \left\{ \begin{matrix} m+1 \\ k \end{matrix} \right\} n^{\underline{k}} - \left\{ \begin{matrix} m+1 \\ m+1 \end{matrix} \right\} n^{\underline{m+1}} - n\sum_{k=1}^{m} \left\{ \begin{matrix} m \\ k-1 \end{matrix} \right\} (n-1)^{\underline{k-1}} \right)\\
=&\frac{1}{n^m}\left( n^{m+1} - n^{\underline{m+1}} - n\sum_{l=0}^{m-1} \left\{ \begin{matrix} m \\ l \end{matrix} \right\} (n-1)^{\underline{l}} \right)\\
=&\frac{1}{n^m}\left( n^{m+1} - n^{\underline{m+1}} - n\sum_{l=1}^{m} \left\{ \begin{matrix} m \\ l \end{matrix} \right\} (n-1)^{\underline{l}} - n\left\{ \begin{matrix} m \\ 0 \end{matrix} \right\} (n-1)^{\underline{0}} + n\left\{ \begin{matrix} m \\ m \end{matrix} \right\} (n-1)^{\underline{m}} \right)\\
=&\frac{1}{n^m}\left( n^{m+1} - n^{\underline{m+1}} - n\left((n-1)^m + 0 - (n-1)^{\underline{m}}\right) \right)\\
=&\frac{1}{n^m}\left( n^{m+1} - n(n-1)^m \right)\\
=&n - n(1-\frac{1}{n})^m\\
=&n\left(1-(1-\frac{1}{n})^m\right)\\
\approx&n\left(1-e^{-\frac{m}{n}}\right) \text{ if n is large enough}
\end{aligned}
\tag{S15}
$$

The binomial distribution of $K$ with probability of $1-\exp(-\frac{m}{n})$ as parameter has an expected value $n(1-\exp(-\frac{m}{n}))$, which is same as the expected value of $k$ conditioned on $m$ as $n$ is large. So there is connection between the conditional distribution of k and the binomial one.

#### 1.2 Multiple Replicates

##### 1.2.1 Probabilistic Framework

With $c$ the concentration of target molecules in the volume, and with $V_d$ the volume to be loaded in the dPCR device, set $\mu = V_d c$, i.e. $\mu$ is the average number of target molecules loaded in the dPCR device. The random sampling (i.e. pipetting) of a volume $V_d$ from the volume will bring a number of target molecules along; this number of molecules is denoted by $M_i$, $i = 1, \ldots, r$, and it is thus considered as a random number. Under ideal conditions it would be reasonable to assume a Poisson distribution, i.e.

$$M_i \sim \text{Poisson}(\mu).$$

See also Fig.1 in the main text. In this setting, the interest is in the estimation of $\mu$ (absolute quantification) or a function of $\mu$ (e.g. CNV= $\mu_A/\mu_B$, with $\mu_A$ and $\mu_B$ the numbers of molecules of type A and B; see further down). Based on a single replicate $i$, the parameter $\mu$ can be estimated as $\frac{V_d}{V_p}\hat{\lambda}_i$ with $\hat{\lambda}_i$ as in Eq.S6. With $r$ replicates, the estimator of $\mu$ can be defined as

$$\hat{\mu} = \frac{V_d}{V_p r} \sum_{i=1}^{r} \hat{\lambda}_i. \tag{S16}$$

The appropriate variance is now $\text{Var}\{\hat{\mu} \mid n_1, \ldots, n_r\}$, which is no longer conditional on the number(s) of molecules, as it must also express the variability over the replicates (replicates involve random sampling from the volume). We will use the shorter notation $\text{Var}\{\hat{\mu} \mid n\}$.

For absolute quantification we find

$$\text{Var}\{\hat{\mu} \mid n\} = \left(\frac{V_d}{V_p r}\right)^2 \sum_{i=1}^{r} \text{Var}\left\{\hat{\lambda}_i \mid n_i\right\}$$

and thus the problem is reduced to finding the variance $\text{Var}\left\{\hat{\lambda}_i \mid n_i\right\}$. As before, since $\hat{\lambda}$ is a function of $K$, we need $\text{Var}\{K_i \mid n_i\}$ and hence the conditional distribution of $K_i \mid n_i$.

The marginal distribution of $K$ is expressed as Eq.1 in the main text. We do not provide an explicit expression for this probability, but an approximation for large numbers of partitions $n_i$, can be obtained. For the detailed proof, see **Section Binomial Distribution of Positive Partitions**. We find,

$$P(K_i = k \mid n_i) \dot{\sim} \frac{n_i!}{(n_i - k)!k!} \left(e^{-\mu/n_i}\right)^{n_i - k} \left(1 - e^{-\mu/n_i}\right)^k, \tag{S17}$$

which is the distribution function of the binomial distribution $\text{Binom}(n_i, \pi_i)$ with $\pi_i = 1 - e^{-\mu/n_i}$ the probability that a partition is positive. Note the difference with the binomial distribution (Eq.S8) for which the probability parameter equals $1 - \exp(-m_i/n_i)$ whereas here we have the marginal mean $\mu$ instead of $m_i$.

##### 1.2.2 The Delta Method for Variance Estimation

The variance $\text{Var}\left\{\hat{\lambda}_i \mid n_i\right\}$ can now be approximated as in **Section Delta Method for the Standard Error** by using (1) the delta method and (2) the variance $\text{Var}\left\{K_i \mid n_i\right\}$ from the binomial distribution (Eq.S17). The unknown parameter $\mu$ must be replaced by its estimator (Eq.S16).

##### 1.2.3 General Approach for Multiplexing

In the next few paragraphs a more generic description is given, which also applies to multiplex experiments. The description is given for experiments with replicates, but at the end it will be indicated how the procedure simplifies when no replicate is available.

For multiplex experiments, for replicate $i = 1, \ldots, r$, let $M_{Ai}, M_{Bi}, \ldots$ denote the randomly sampled numbers of target molecules of type A, B, ... that are partitioned over the $n_i$ partitions. Let $\boldsymbol{M}_i^t = (M_{Ai}, M_{Bi}, \ldots)$, $\boldsymbol{M}^t = (\boldsymbol{M}_1^t, \ldots, \boldsymbol{M}_r^t)$ and $\boldsymbol{n}^t = (n_1, \ldots, n_r)$. The numbers of positive partitions for types of targets A, B, ... in replicate $i$ are denoted by $K_{Ai}, K_{Bi}, \ldots$. Let $\boldsymbol{K}_i^t = (K_{Ai}, K_{Bi}, \ldots)$ and $\boldsymbol{K}^t = (\boldsymbol{K}_1^t, \ldots, \boldsymbol{K}_r^t)$. For all types of target molecules, let $\mu_A, \mu_B, \ldots$ denote the average numbers of molecules in a fixed volume $V_d$ loaded in the dPCR device. The parameters $\lambda_{Ai}, \lambda_{Bi}, \ldots$ refer to the average numbers of molecules of type A, B, ... per partition in replicate $i$.

Suppose that the goal of the experiment is to estimate a parameter $\theta$ which can be expressed as a function of the $\mu$ (or $\lambda$) parameters. An estimator of $\theta$ can be obtained by replacing all $\mu$ (or $\lambda$) parameters by their estimators, which in turn depend on $\boldsymbol{K}$ and $\boldsymbol{n}$. It will be convenient to also explicitly consider the estimator $\hat{\theta}$ as a function of $\boldsymbol{M}$, because the distribution of $\boldsymbol{K}$ depends on it. We therefore write the estimator of $\theta$ as $\hat{\theta} = \hat{\theta}(\boldsymbol{M}, \boldsymbol{K}, \boldsymbol{n})$. We are now interested in the estimation of its variance, $\text{Var}_{MX}\left\{\hat{\theta} \mid \boldsymbol{n}\right\}$, which is estimated as the empirical variance over the replicates and covers both sampling and random partitioning variability.

In the special case of no replicates (i.e. $r = 1$), we cannot make a distinction between the concentration of target molecules in the vessel and the concentration of the target loaded on the dPCR device. Hence, $\lambda$ takes over the role of $\mu$, and the number of target molecules is considered fixed, i.e. we use $m$ instead of $M$. This number is no longer considered random and so we only need the conditional variance $\mathrm{Var}_{K|M}\left\{\hat{\theta} \mid \boldsymbol{M} = \boldsymbol{m}, \boldsymbol{n}\right\}$, which follows immediately from the conditional bootstrap approach of **Section A Conditional Bootstrap Method for Standard Errors** in the main text.

### 2 Simulation Settings

We simulate the process for several orders of magnitude of concentration reflecting empirical dilution levels. Therefore we vary the expected number of target molecules per partition $\lambda$ from 0.005 to 1.5. The pipetting error we added is normally distributed with a coefficient of variation of 3%. The number of partitions is initially set at 20 000. Partitions are assumed to be lost completely at random. To simulate this process, we randomly retain partitions between replicates with an expected value 16000 and standard deviation 2000. Then the partition size is modeled to follow a log-normal distribution with mean 0 and standard deviation 0.1, which is approximately equal to a normal distribution with a coefficient of variation of 10%. In the final stage, partitions are classifiied as positive or negative after thresholding. To assess the effect of misclassification, we set 5% false negative rate and 0.01% false positive rate. Those simulation studies are implemented in parallel (separately) and sequentially.

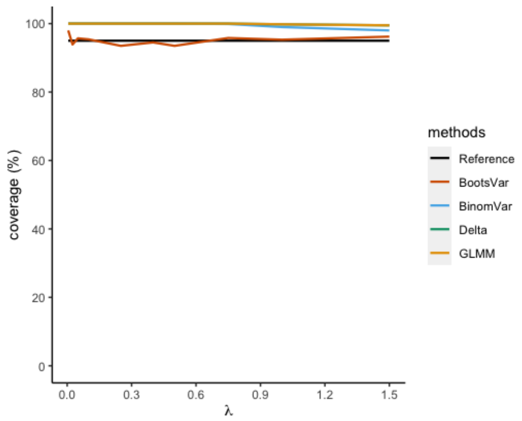

Figure S1: Empirical coverage of 95% CIs in absolute quantification with a single replicate given a fixed $m$ and $n$

### 3 Conditional Variance

| **lambda ($\lambda$)** | **BootsVar** | **BinomVar** | **Delta** | **GLMM** |
|---|---|---|---|---|
| 0.00555 | 64.70 | 35416.81 | 35299.45 | 35299.40 |
| 0.02355 | 6.82 | 8628.35 | 8630.73 | 8629.30 |
| 0.051 | -1.14 | 3781.38 | 3780.61 | 3780.61 |
| 0.10365 | -1.35 | 1826.21 | 1826.35 | 1826.30 |
| 0.25095 | -9.44 | 652.90 | 653.03 | 653.03 |
| 0.3978 | -6.29 | 403.98 | 403.76 | 403.76 |
| 0.5092 | -6.28 | 300.98 | 301.36 | 301.36 |
| 0.74825 | 7.09 | 227.14 | 226.67 | 226.67 |
| 0.9949 | 4.51 | 150.48 | 150.53 | 150.53 |
| 1.4922 | 2.38 | 80.21 | 80.16 | 80.16 |

Table S1: Relative Bias (%) of several estimators of the conditional variance with a single replicate (given a fixed m and n).

| Methods | Runtime (s) |
|---|---|
| **BootsVar** | 4.0386 |
| **BinomVar** | 0.0146 |
| **GLMM** | 0.1675 |
| **Delta Method** | 0.0133 |

Table S2: Computation time (4GB 1600 MHz DDR3) for 1 simulation run with 80% of partitions positive and no replicates. For BootsVar and BinomVar, the bootstrap iteration is set at 1000.

### 4 Absolute Quantification

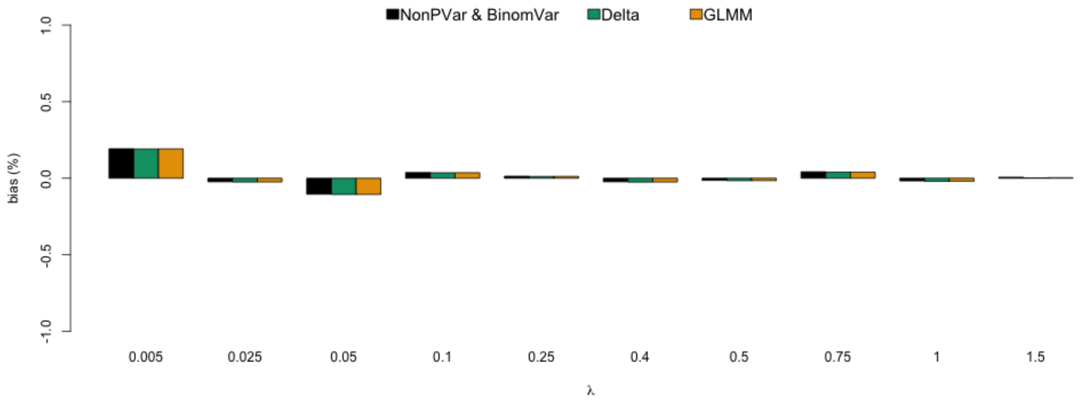

Figure S2: Relative bias of the estimators of $\lambda$ in absolute quantification (3 replicates) with only sampling variation and random partitioning.

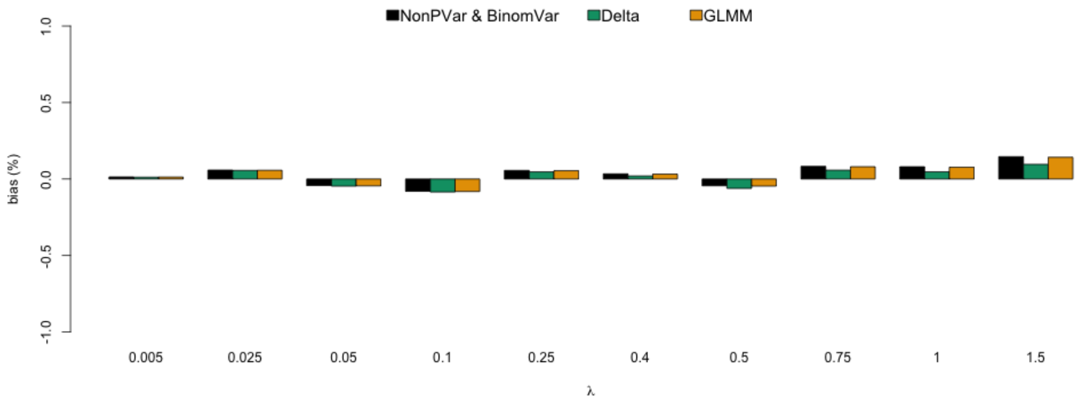

Figure S3: Relative bias of the estimator of $\lambda$ in absolute quantification (3 replicates) with additional pipetting error.

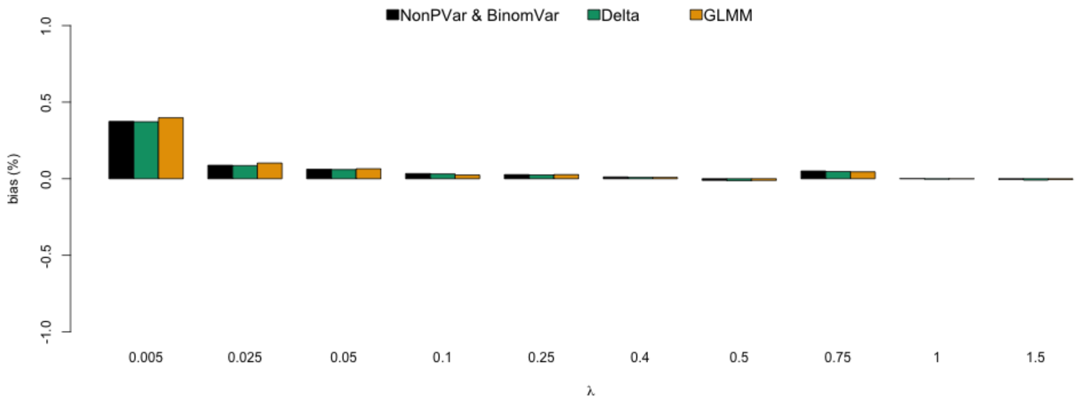

Figure S4: Relative bias of the estimator of $\lambda$ in absolute quantification (3 replicates) with additional partition loss.

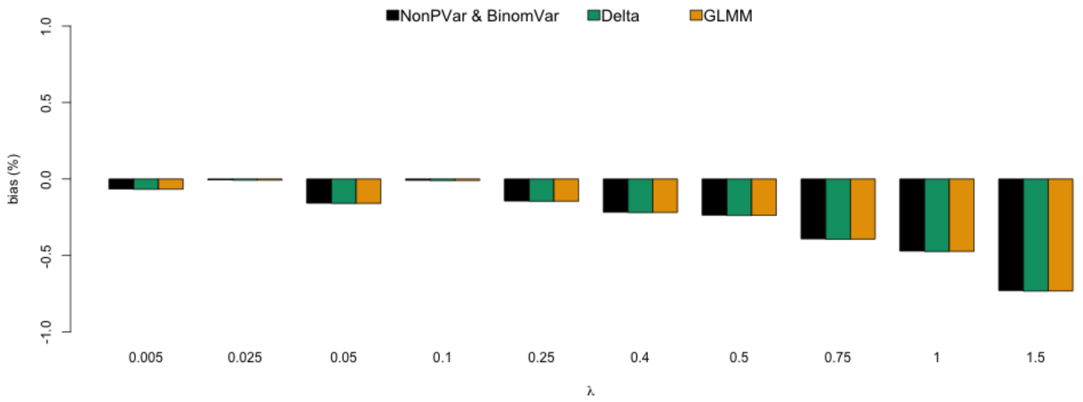

Figure S5: Relative bias of the estimator of $\lambda$ in absolute quantification (3 replicates) with additional partition size variation.

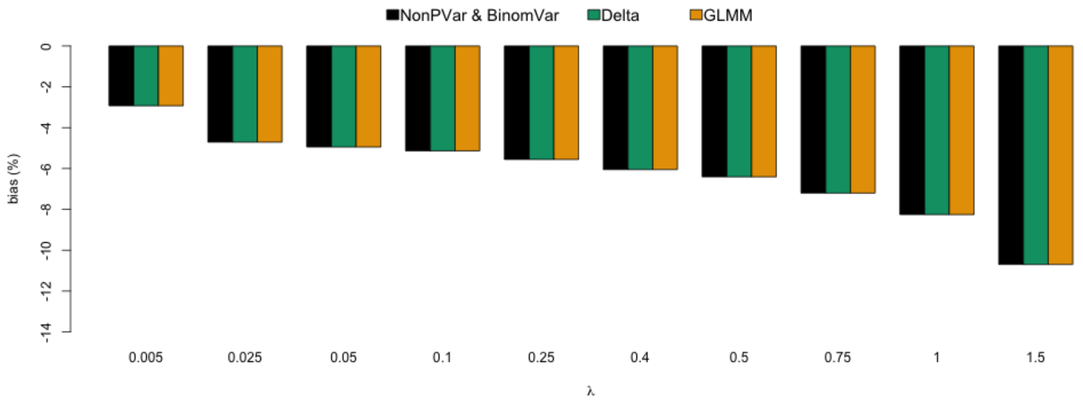

Figure S6: Relative bias of the estimator of $\lambda$ in absolute quantification (3 replicates) with additional misclassification.

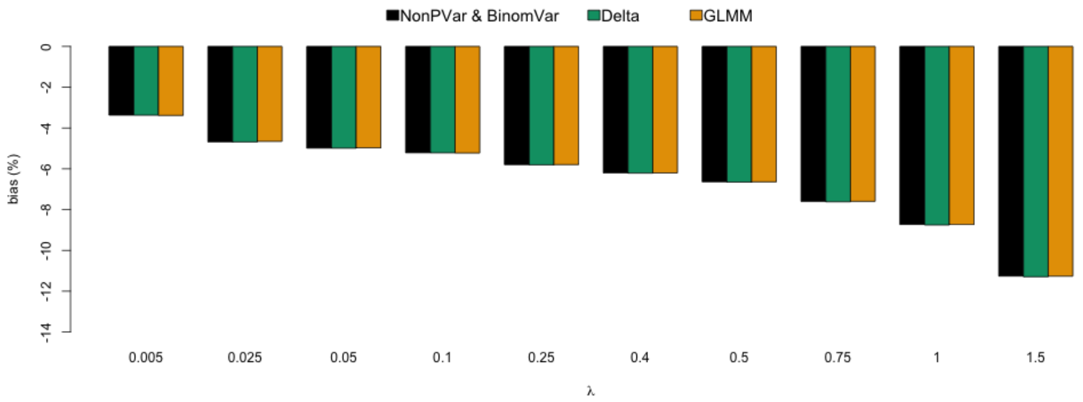

Figure S7: Relative bias of the estimator of $\lambda$ in absolute quantification (3 replicates) with all sources of variation.

The absolute bias is calculated as,

$$\text{bias}(log) = \log\left(|Var(\hat{\lambda}) - Var(\lambda)|\right)$$

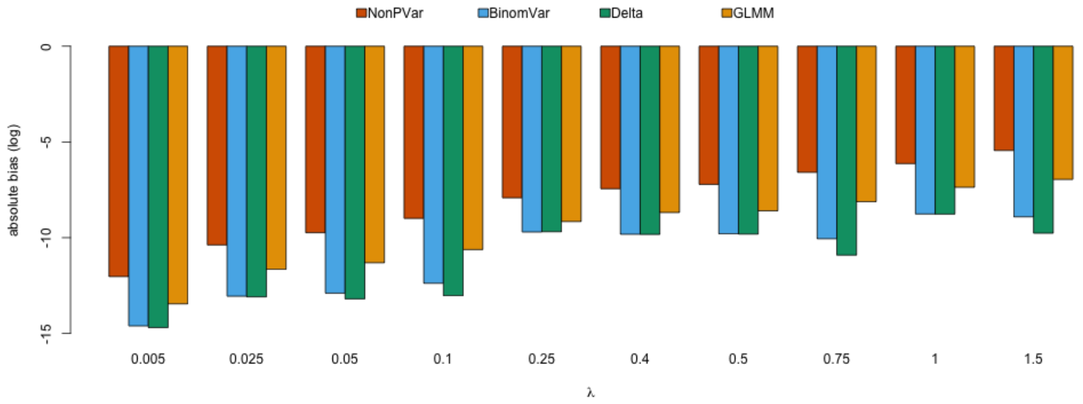

Figure S8: Absolute bias of the variance estimator of $\lambda$ in absolute quantification (3 replicates) with only sampling variation and random partitioning. Note the absolute bias is on the log scale. Smaller on the log scale means closer to 0 on the original scale.

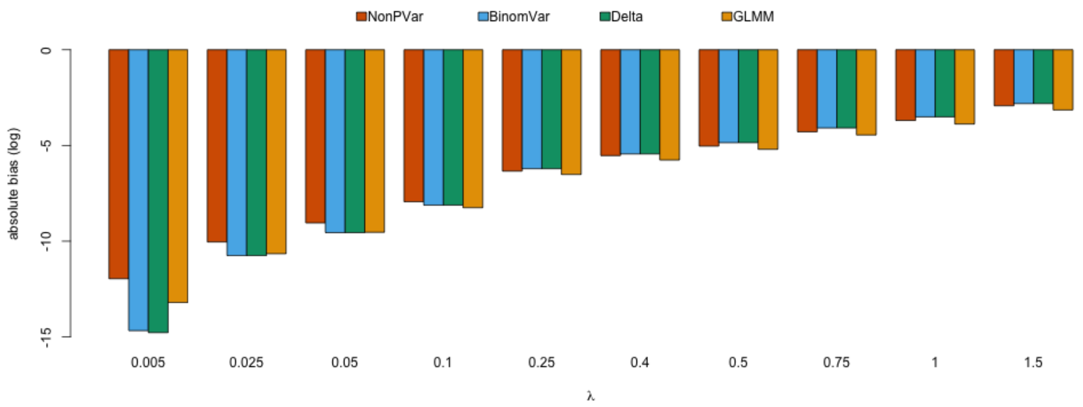

Figure S9: Absolute bias of the variance estimator of $\lambda$ in absolute quantification (3 replicates) with additional pipetting error. Note the absolute bias is on the log scale. Smaller on the log scale means closer to 0 on the original scale.

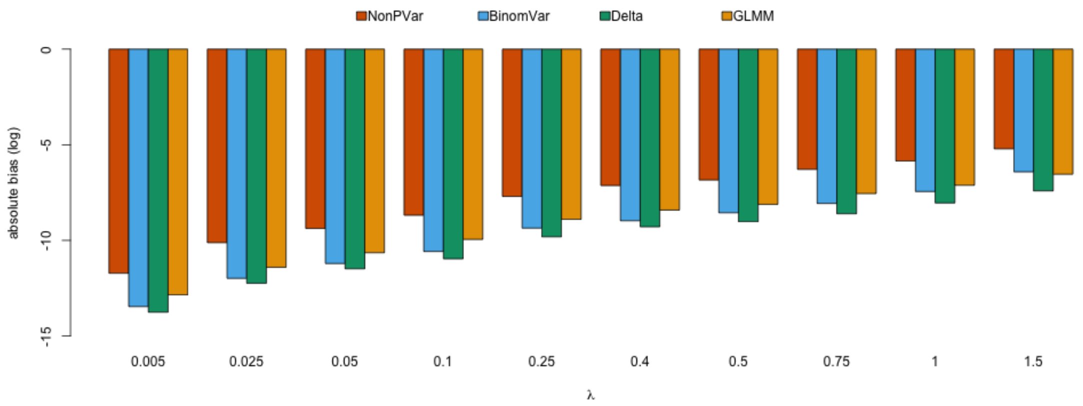

Figure S10: Absolute bias of the variance estimator of $\lambda$ in absolute quantification (3 replicates) with additional partition loss. Note the absolute bias is on the log scale. Smaller on the log scale means closer to 0 on the original scale.

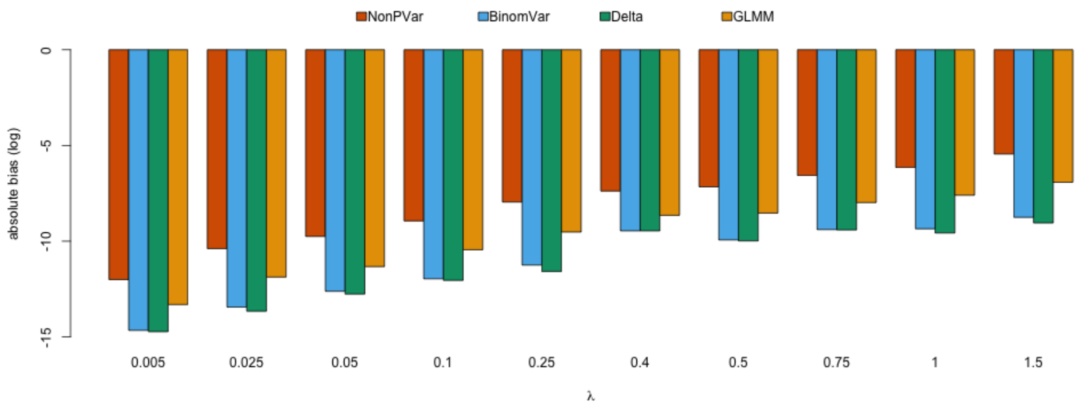

Figure S11: Absolute bias of the variance estimator of $\lambda$ in absolute quantification (3 replicates) with additional partition size variation. Note the absolute bias is on the log scale. Smaller on the log scale means closer to 0 on the original scale.

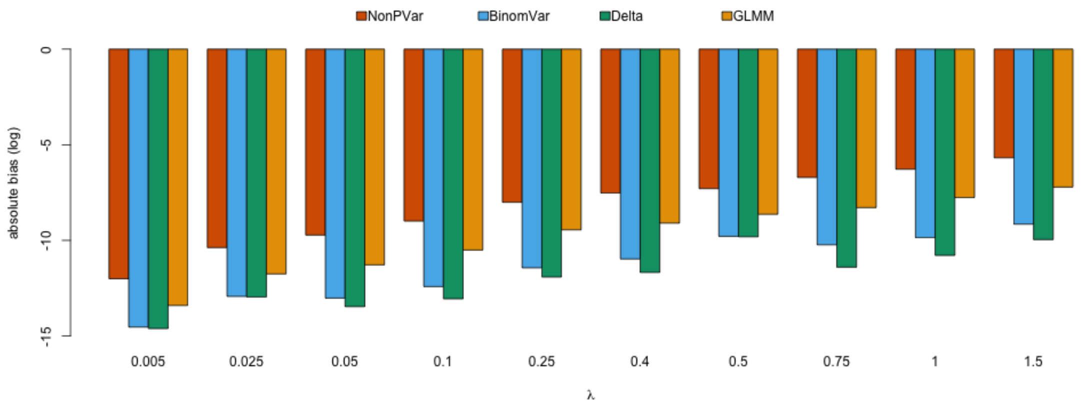

Figure S12: Absolute bias of the variance estimator of $\lambda$ in absolute quantification (3 replicates) with additional misclassification. Note the absolute bias is on the log scale. Smaller on the log scale means closer to 0 on the original scale.

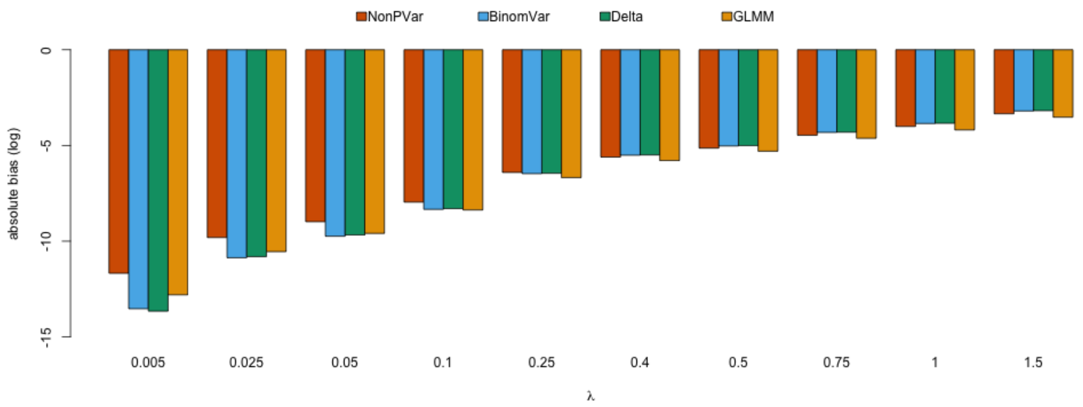

Figure S13: Absolute bias of the variance estimator of $\lambda$ in absolute quantification (3 replicates) with all sources of variation. Note the absolute bias is on the log scale. Smaller on the log scale means closer to 0 on the original scale.

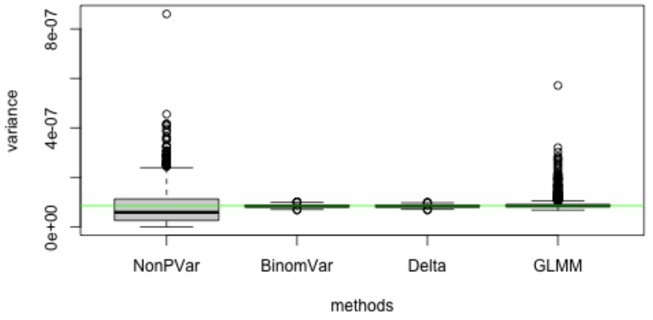

Figure S14: Variance estimates of simulation runs in absolute quantification (3 replicates) with only sampling variation and random partitioning in low concentration setting of $\lambda_A = 0.005$. The horizontal line is the estimated variance in the simulation (a good approximation to the true variance).

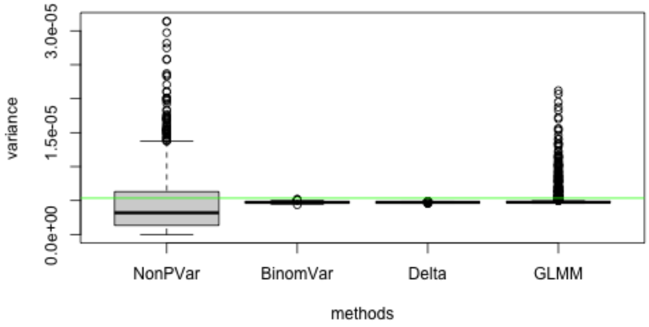

Figure S15: Variance estimates of simulation runs in absolute quantification (3 replicates) with only sampling variation and random partitioning in medium concentration setting of $\lambda_A = 0.25$. The horizontal line is the estimated variance in the simulation (a good approximation to the true variance).

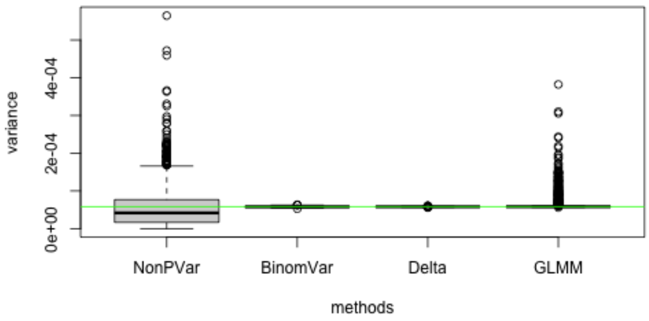

Figure S16: Variance estimates of simulation runs in absolute quantification (3 replicates) with only sampling variation and random partitioning in high concentration setting of $\lambda_A = 1.5$. The horizontal line is the estimated variance in the simulation (a good approximation to the true variance).

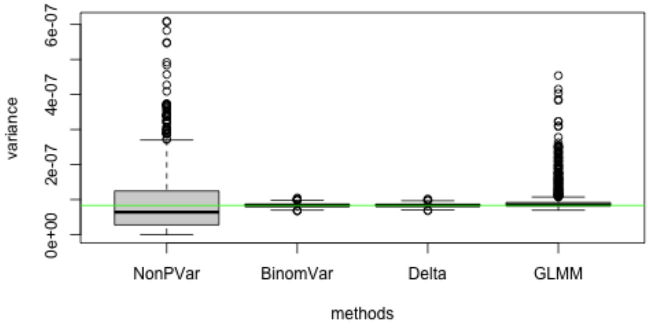

Figure S17: Variance estimates of simulation runs in absolute quantification (3 replicates) with additional pipetting error in low concentration setting of $\lambda_A = 0.005$. The horizontal line is the estimated variance in the simulation (a good approximation to the true variance).

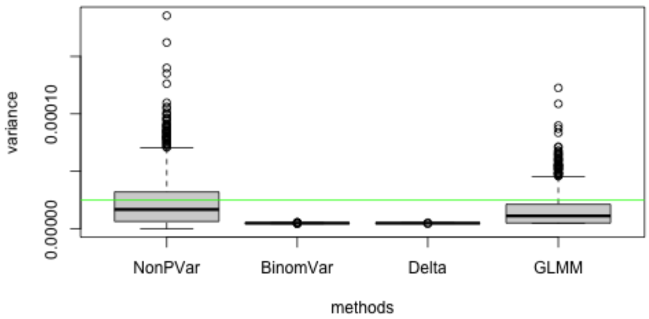

Figure S18: Variance estimates of simulation runs in absolute quantification (3 replicates) with additional pipetting error in medium concentration setting of $\lambda_A = 0.25$. The horizontal line is the estimated variance in the simulation (a good approximation to the true variance).

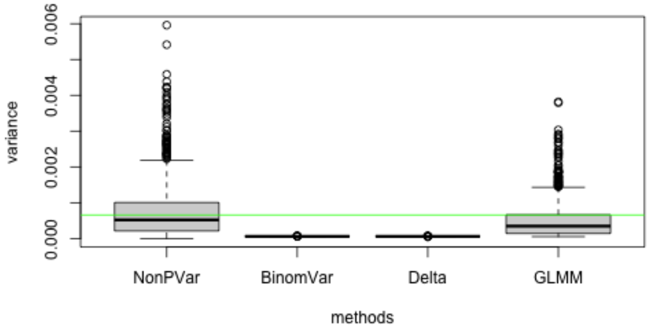

Figure S19: Variance estimates of simulation runs in absolute quantification (3 replicates) with additional pipetting error in high concentration setting of $\lambda_A = 1.5$. The horizontal line is the estimated variance in the simulation (a good approximation to the true variance).

### 5 Copy Number Variation

#### 5.1 Singleplex

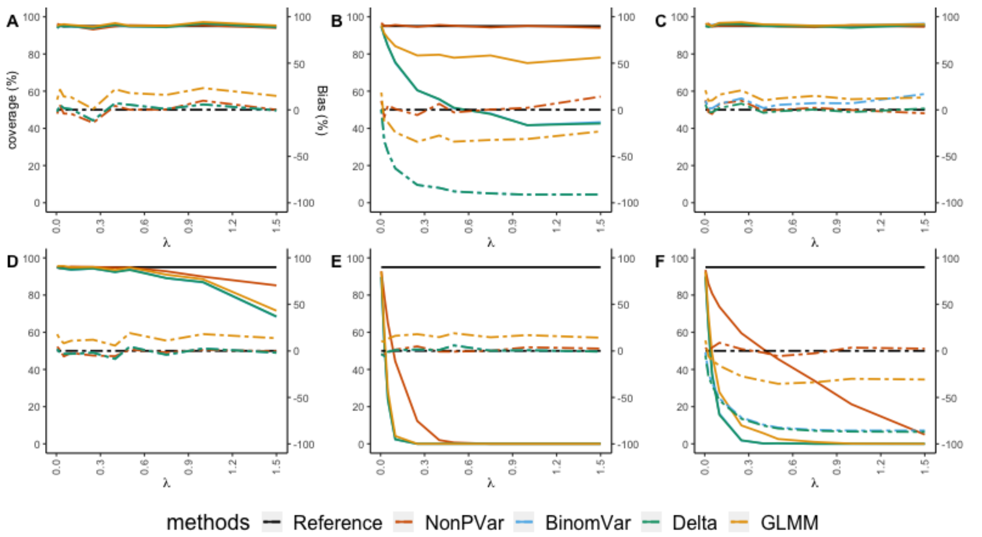

Figure S20: Empirical coverage of the 95% CIs (solid lines, left axis) and relative bias (dashed lines, right axis) for CNV in singleplex in different scenarios. X-axis represents varying concentration of target molecules from low to high. (A) only sampling variation and random partitioning with 3 replicates (B) 3% pipetting error (C) 20% partition loss (D) coefficient of variation of 10% in partition size (E) misclassification with 0.01% false positive rate and 5% false negative rate (F) all variation included. The reference (black solid line) for empirical coverage is set at 95%. The constructed CIs are supposed to cover the true values in 95% of the time. The closer other solid lines are to this reference, the better the CIs are. The reference (black dashed line) is set at 0%. The closer other dashed lines are to this reference, the lower the relative bias is.

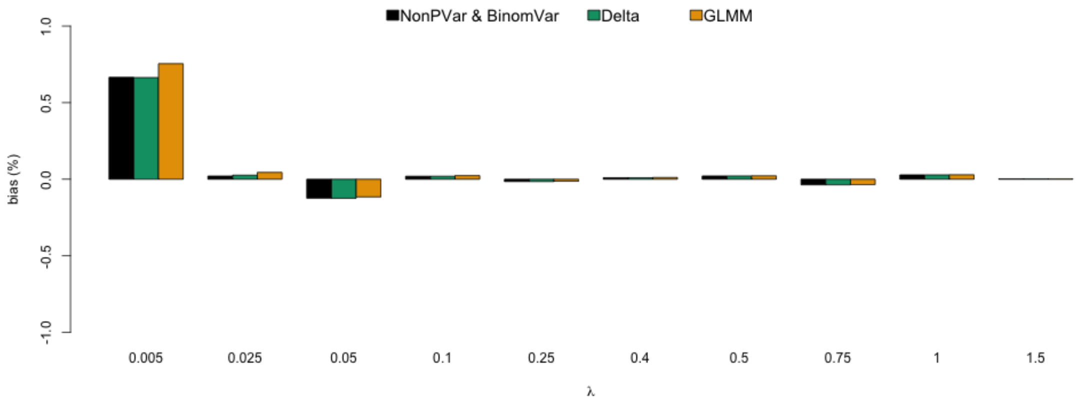

Figure S21: Relative bias of the estimator of cnv in CNV singleplex (3 replicates) with only sampling variation and random partitioning.

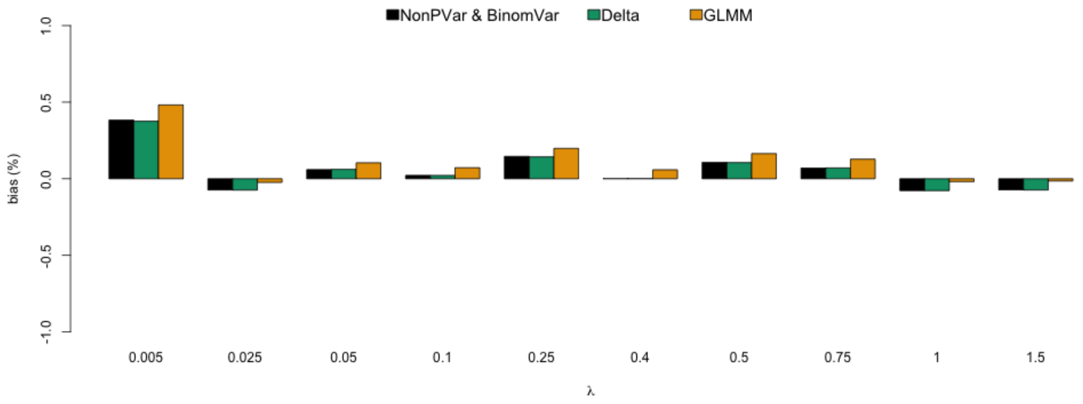

Figure S22: Relative bias of the estimator of cnv in CNV singleplex (3 replicates) with additional pipetting error.

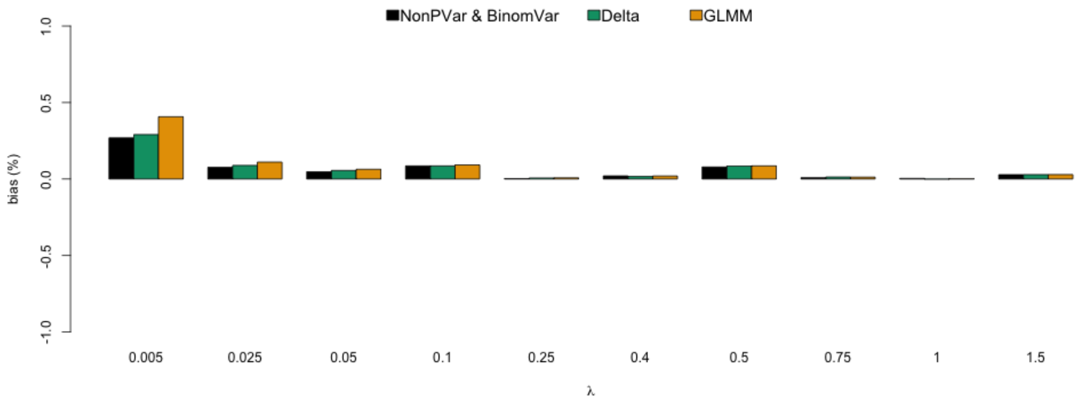

Figure S23: Relative bias of the estimator of cnv in CNV singleplex (3 replicates) with additional partition loss.

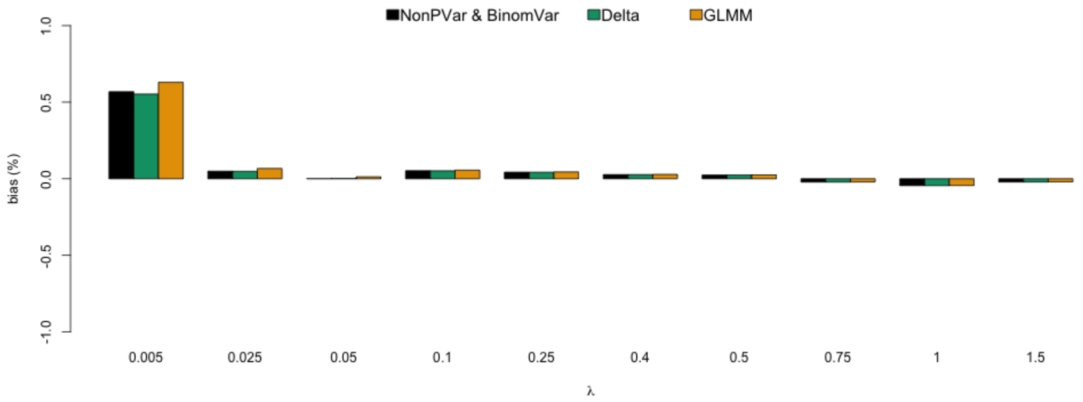

Figure S24: Relative bias of the estimator of cnv in CNV singleplex (3 replicates) with additional partition size variation.

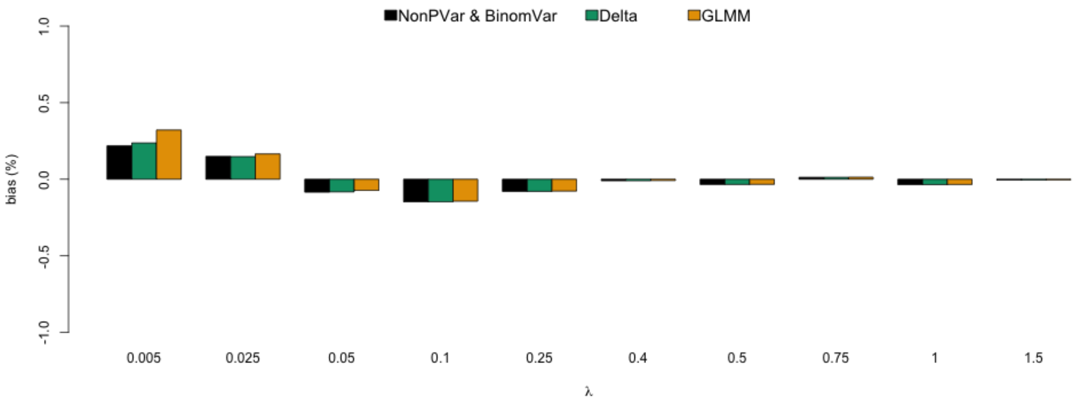

Figure S25: Relative bias of the estimator of cnv in CNV singleplex (3 replicates) with additional misclassification.

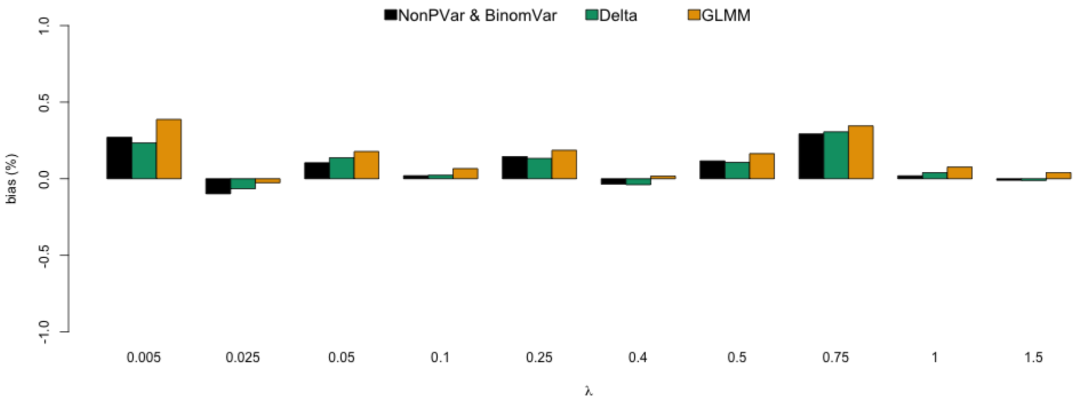

Figure S26: Relative bias of the estimator of cnv in CNV singleplex (3 replicates) with all sources of variation.

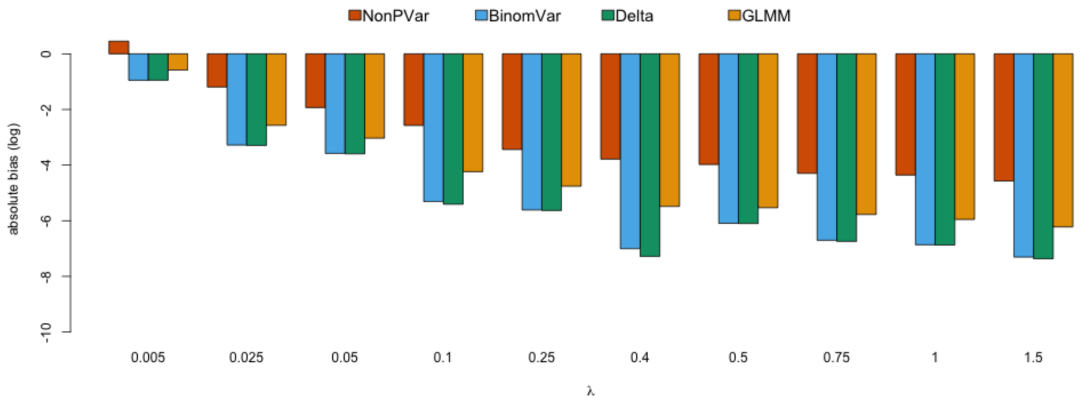

Figure S27: Absolute bias of the variance estimator of cnv in CNV singleplex (3 replicates) with only sampling variation and random partitioning.Note the absolute bias is on the log scale. Smaller on the log scale means closer to 0 on the original scale.

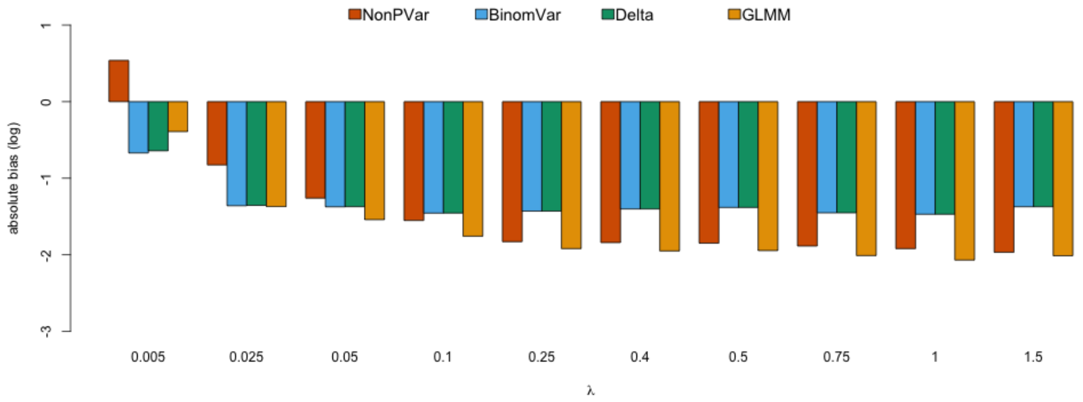

Figure S28: Absolute bias of the variance estimator of cnv in CNV singleplex (3 replicates) with additional pipetting error. Note the absolute bias is on the log scale. Smaller on the log scale means closer to 0 on the original scale.

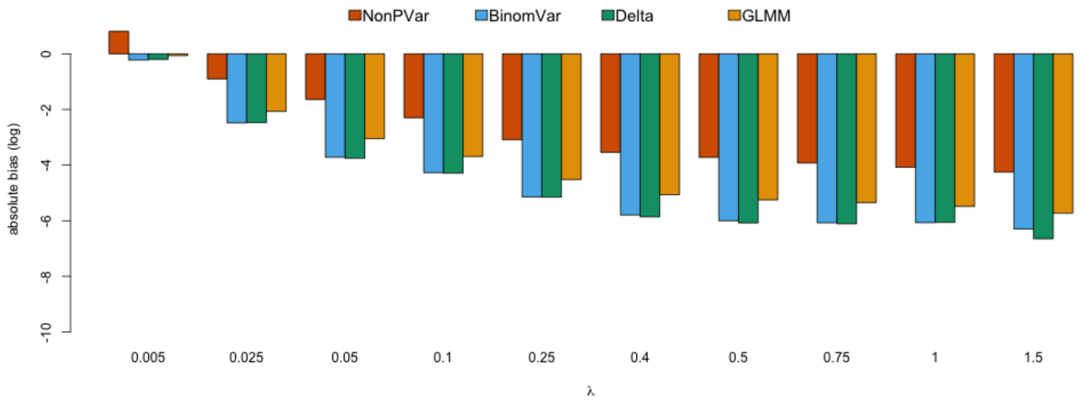

Figure S29: Absolute bias of the variance estimator of cnv in CNV singleplex (3 replicates) with additional partition loss. Note the absolute bias is on the log scale. Smaller on the log scale means closer to 0 on the original scale.

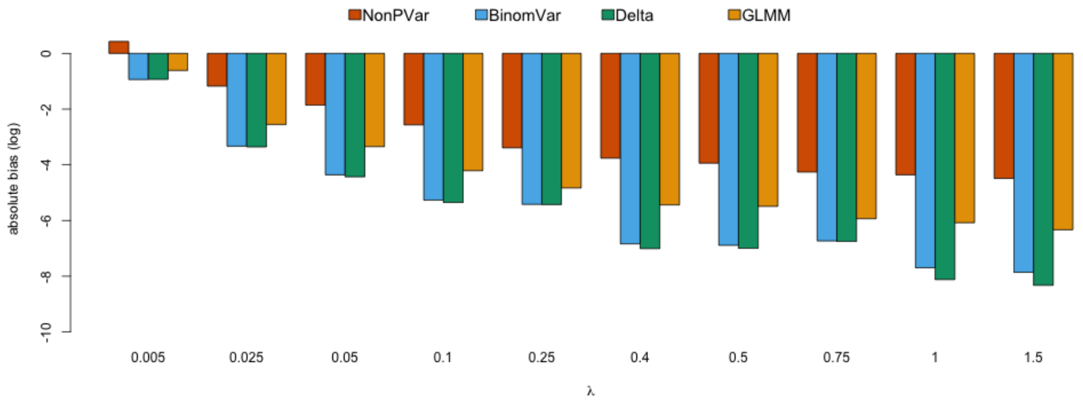

Figure S30: Absolute bias of the variance estimator of cnv in CNV singleplex (3 replicates) with additional partition size variation. Note the absolute bias is on the log scale. Smaller on the log scale means closer to 0 on the original scale.

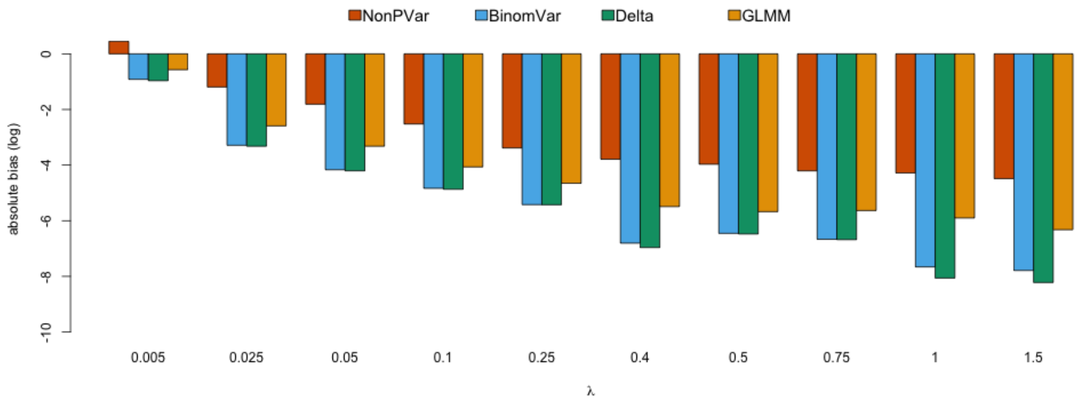

Figure S31: Absolute bias of the variance estimator of cnv in CNV singleplex (3 replicates) with additional misclassification. Note the absolute bias is on the log scale. Smaller on the log scale means closer to 0 on the original scale.

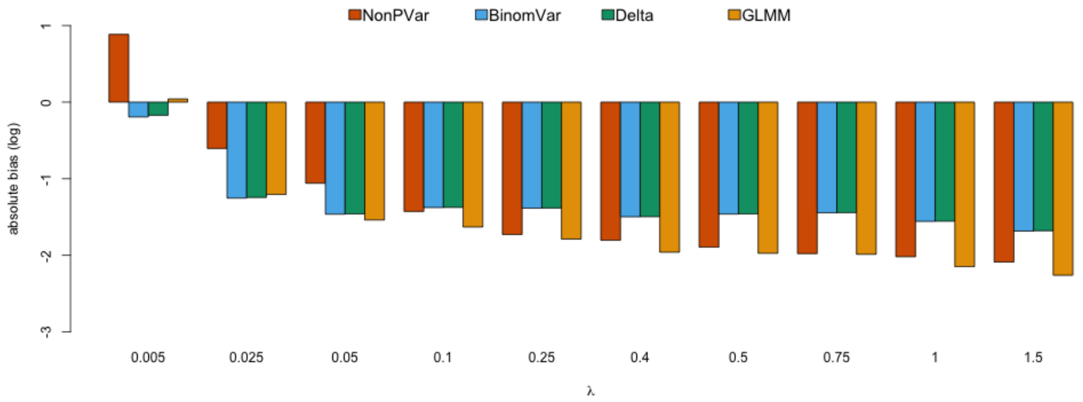

Figure S32: Absolute bias of the variance estimator of cnv in CNV singleplex (3 replicates) with all sources of variation. Note the absolute bias is on the log scale. Smaller on the log scale means closer to 0 on the original scale.

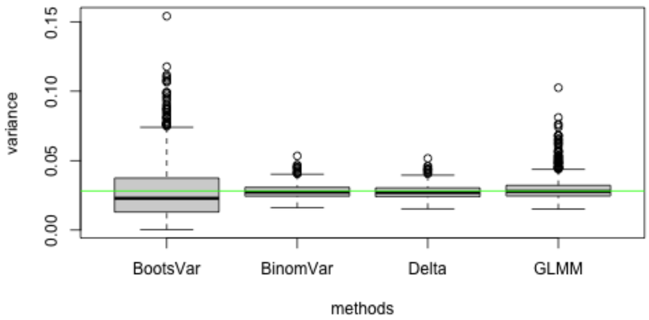

Figure S33: Variance estimates of simulation runs in CNV singleplex (3 replicates) with only sampling variation and random partitioning in low concentration setting of $\lambda_A = 0.005$. The horizontal line is the estimated variance in the simulation (a good approximation to the true variance).

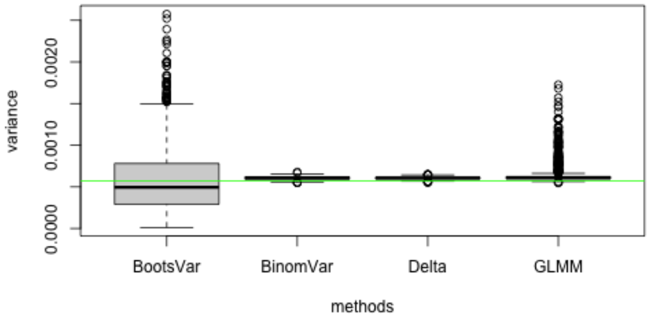

Figure S34: Variance estimates of simulation runs in CNV singleplex (3 replicates) with only sampling variation and random partitioning in medium concentration setting of $\lambda_A = 0.25$. The horizontal line is the estimated variance in the simulation (a good approximation to the true variance).

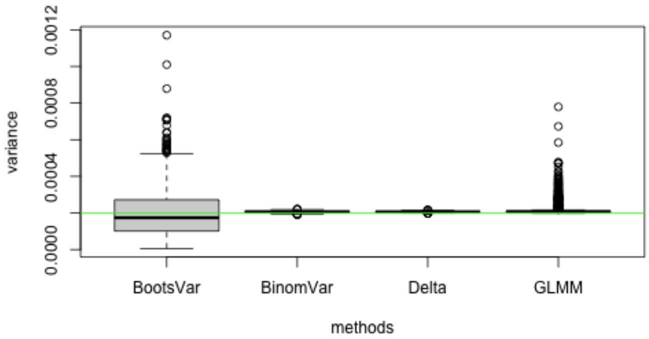

Figure S35: Variance estimates of simulation runs in CNV singleplex (3 replicates) with only sampling variation and random partitioning in high concentration setting of $\lambda_A = 1.5$. The horizontal line is the estimated variance in the simulation (a good approximation to the true variance).

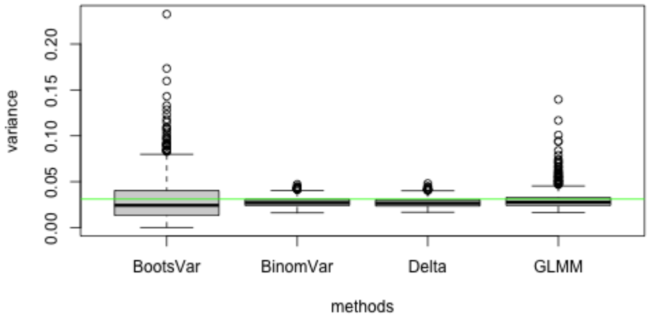

Figure S36: Variance estimates of simulation runs in CNV singleplex (3 replicates) with additional pipetting error in low concentration setting of $\lambda_A = 0.005$. The horizontal line is the estimated variance in the simulation (a good approximation to the true variance).

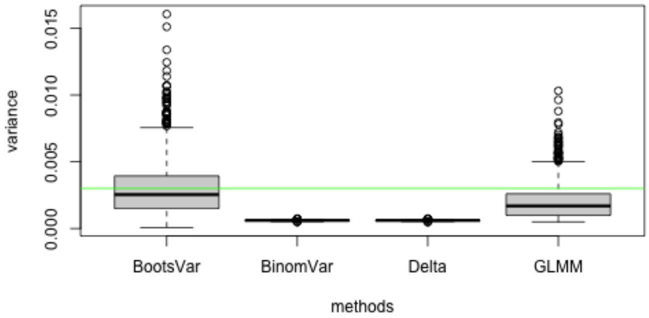

Figure S37: Variance estimates of simulation runs in CNV singleplex (3 replicates) with additional pipetting error in medium concentration setting of $\lambda_A = 0.25$. The horizontal line is the estimated variance in the simulation (a good approximation to the true variance).

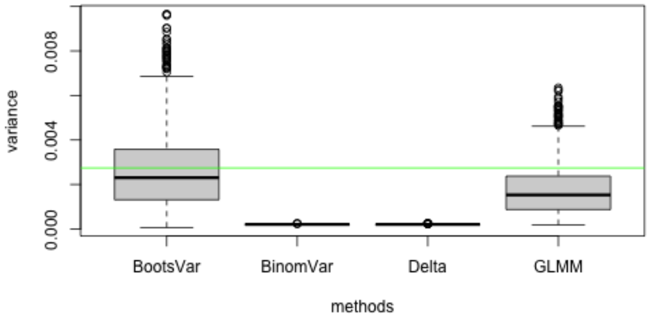

Figure S38: Variance estimates of simulation runs in CNV singleplex (3 replicates) with additional pipetting error in high concentration setting of $\lambda_A = 1.5$. The horizontal line is the estimated variance in the simulation (a good approximation to the true variance).

#### 5.2 Duplex

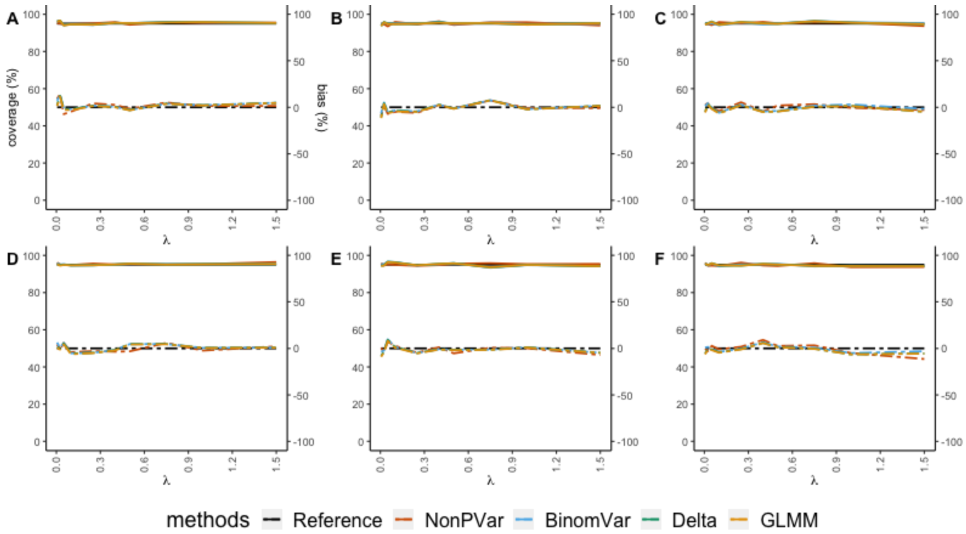

Figure S39: Empirical coverage of the 95% CIs (solid lines, left axis) and relative bias (dashed lines, right axis) for CNV in duplex in different scenarios. X-axis represents varying concentration of target molecules from low to high. (A) only sampling variation and random partitioning with 3 replicates (B) 3% pipetting error (C) 20% partition loss (D) coefficient of variation of 10% in partition size (E) misclassification with 0.01% false positive rate and 5% false negative rate (F) all variation included. The reference (black solid line) for empirical coverage is set at 95%. The constructed CIs are supposed to cover the true values in 95% of the time. The closer other solid lines are to this reference, the better the CIs are. The reference (black dashed line) is set at 0%. The closer other dashed lines are to this reference, the lower the relative bias is.

### 6 Mutational Load

#### 6.1 Low Mutational Load

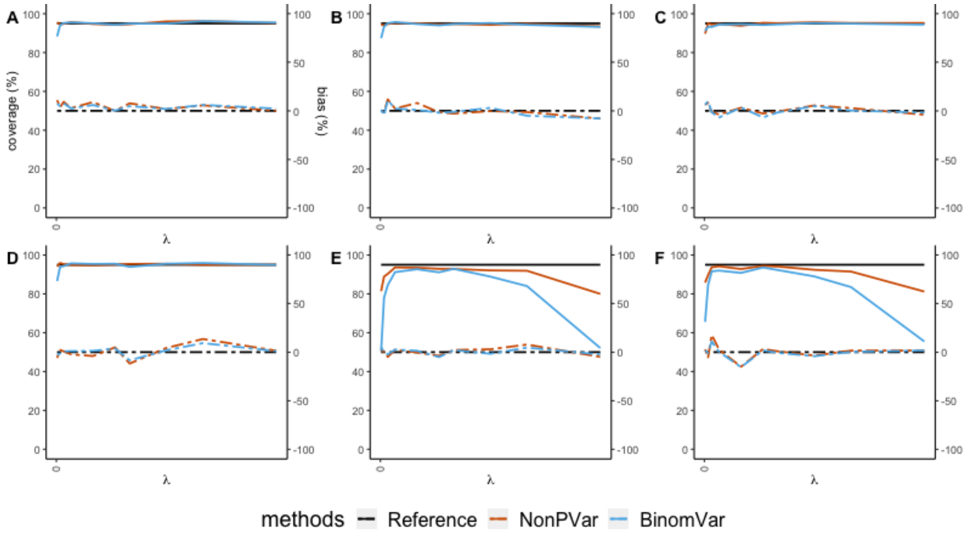

Figure S40: Empirical coverage of the 95% CIs (solid lines, left axis) and relative bias (dashed lines, right axis) for for low mutational load (=1%) in different scenarios. X-axis represents varying concentration of mutant type from low to high. (A) only sampling variation and random partitioning with 3 replicates (B) 3% pipetting error (C) 20% partition loss (D) coefficient of variation of 10% in partition size (E) misclassification with 0.01% false positive rate and 5% false negative rate (F) all variation included. The reference (black solid line) for empirical coverage is set at 95%. The constructed CIs are supposed to cover the true values in 95% of the time. The closer other solid lines are to this reference, the better the CIs are. The reference (black dashed line) is set at 0%. The closer other dashed lines are to this reference, the lower the relative bias is.

#### 6.2 Medium Mutational Load

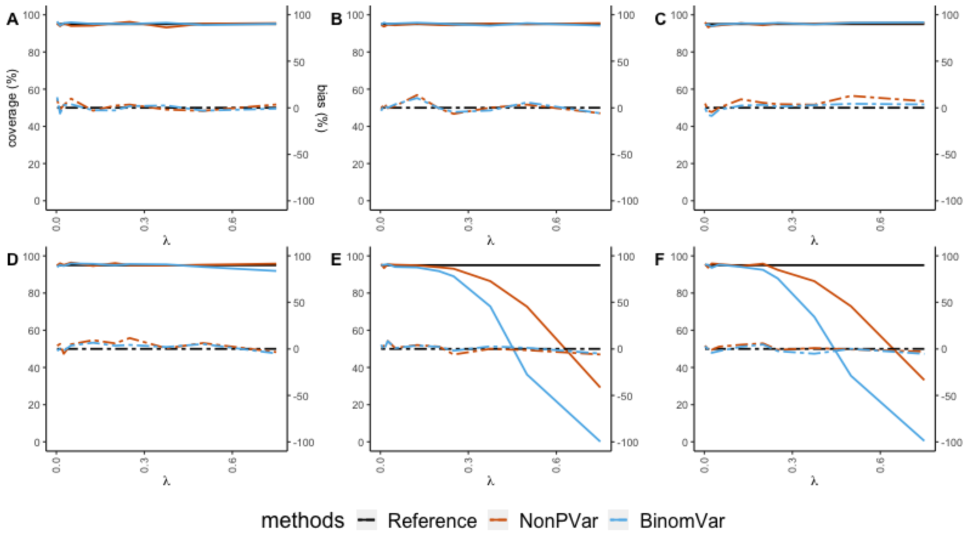

Figure S41: Empirical coverage of the 95% CIs (solid lines, left axis) and relative bias (dashed lines, right axis) for for medium mutational load (=33%) in different scenarios. X-axis represents varying concentration of mutant type from low to high. (A) only sampling variation and random partitioning with 3 replicates (B) 3% pipetting error (C) 20% partition loss (D) coefficient of variation of 10% in partition size (E) misclassification with 0.01% false positive rate and 5% false negative rate (F) all variation included. The reference (black solid line) for empirical coverage is set at 95%. The constructed CIs are supposed to cover the true values in 95% of the time. The closer other solid lines are to this reference, the better the CIs are. The reference (black dashed line) is set at 0%. The closer other dashed lines are to this reference, the lower the relative bias is.

#### 6.3 High Mutational Load

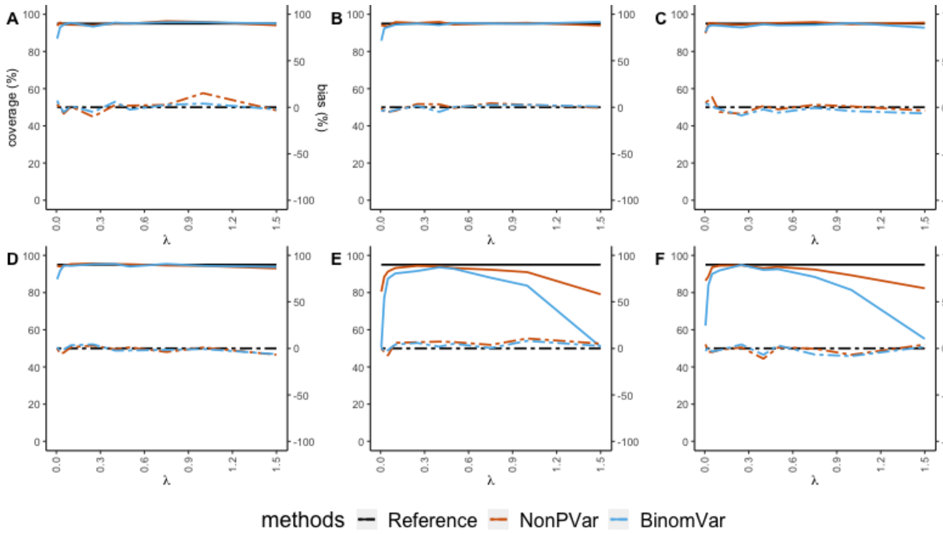

Figure S42: Empirical coverage of the 95% CIs (solid lines, left axis) and relative bias (dashed lines, right axis) for for high mutational load (=99%) in different scenarios. X-axis represents varying concentration of mutant type from low to high. (A) only sampling variation and random partitioning with 3 replicates (B) 3% pipetting error (C) 20% partition loss (D) coefficient of variation of 10% in partition size (E) misclassification with 0.01% false positive rate and 5% false negative rate (F) all variation included. The reference (black solid line) for empirical coverage is set at 95%. The constructed CIs are supposed to cover the true values in 95% of the time. The closer other solid lines are to this reference, the better the CIs are. The reference (black dashed line) is set at 0%. The closer other dashed lines are to this reference, the lower the relative bias is.

### 7 DNA Shearing Index

#### 7.1 Low DSI

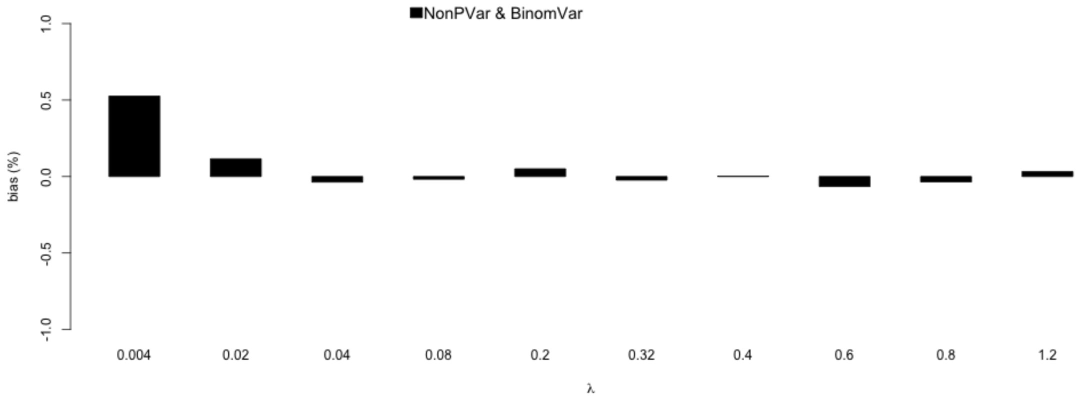

Figure S43: Relative bias of the estimator of dsi in low DSI (=20%, 3 replicates) multiplex with only sampling variation and random partitioning.

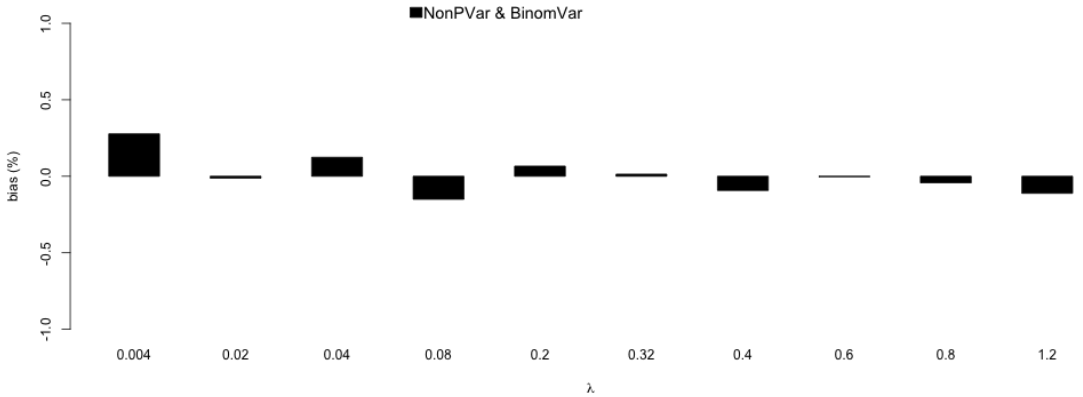

Figure S44: Relative bias of the estimator of dsi in low DSI (=20%, 3 replicates) multiplex with additional pipetting error.

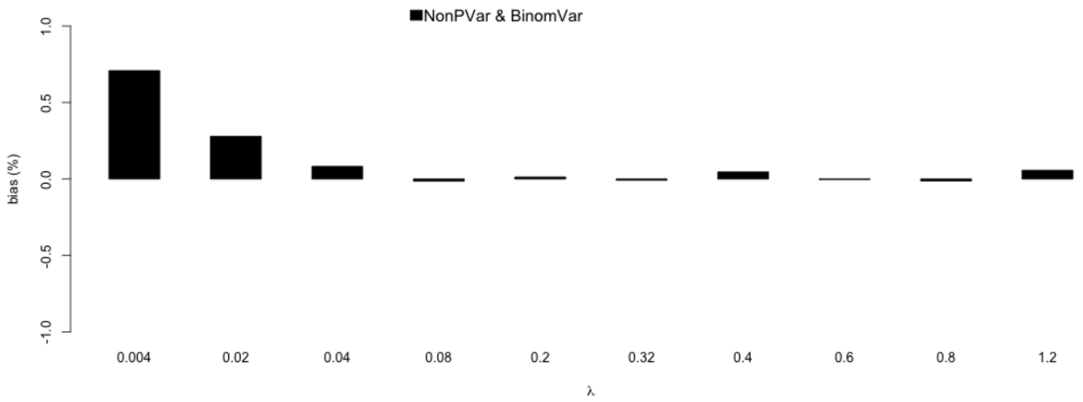

Figure S45: Relative bias of the estimator of dsi in low DSI (=20%, 3 replicates) multiplex with additional partition loss.

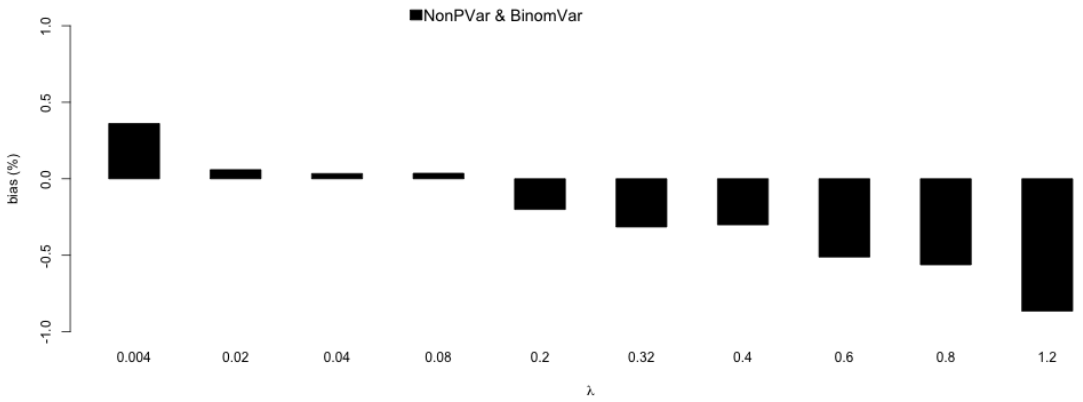

Figure S46: Relative bias of the estimator of dsi in low DSI (=20%, 3 replicates) multiplex with additional partition size variation.

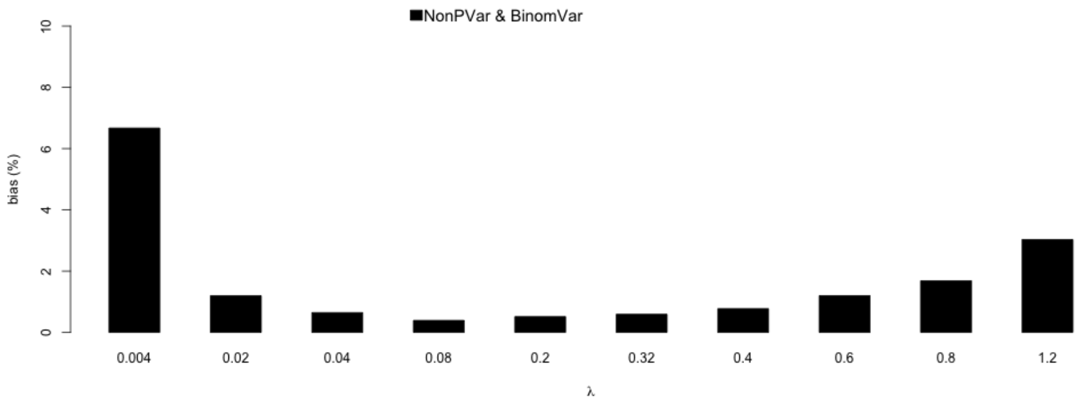

Figure S47: Relative bias of the estimator of dsi in low DSI (=20%, 3 replicates) multiplex with additional misclassification.

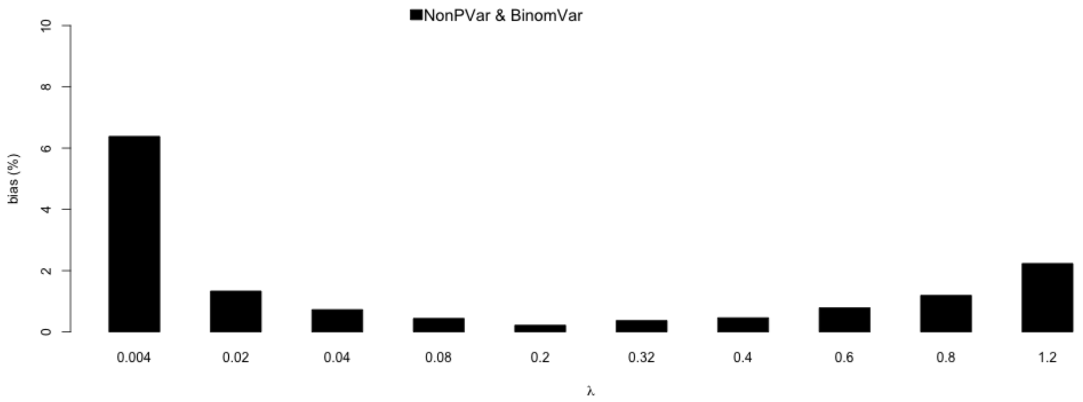

Figure S48: Relative bias of the estimator of dsi in low DSI (=20%, 3 replicates) multiplex with all sources of variation.

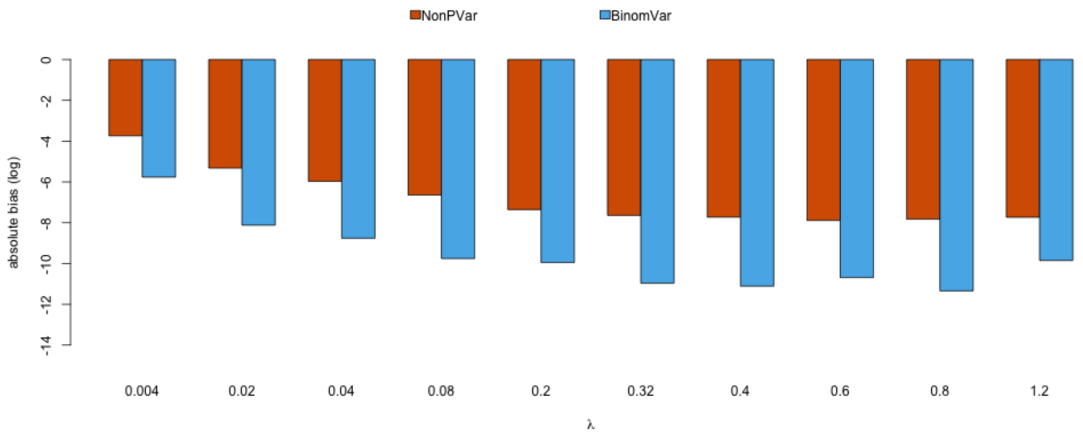

Figure S49: Absolute bias of the variance estimator of dsi in low DSI (=20%, 3 replicates) multiplex with only sampling variation and random partitioning. Note the absolute bias is on the log scale. Smaller on the log scale means closer to 0 on the original scale.

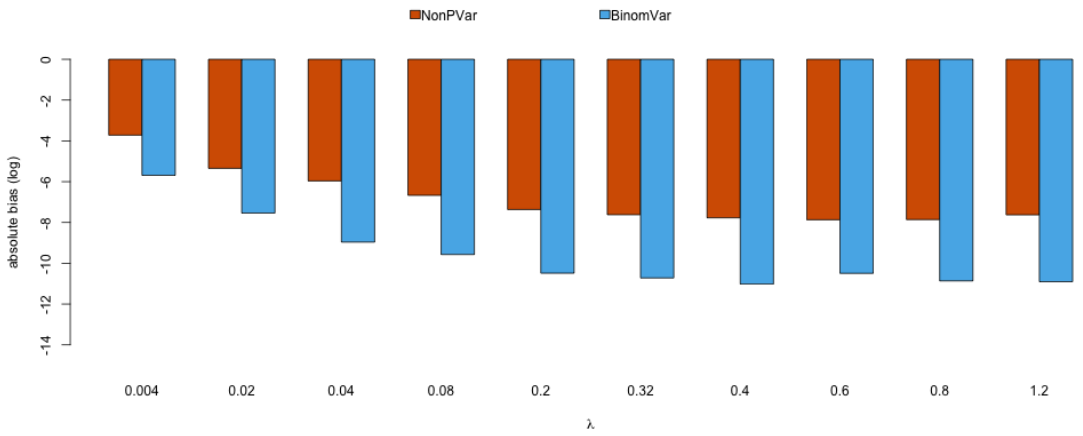

Figure S50: Absolute bias of the variance estimator of dsi in low DSI (=20%, 3 replicates) multiplex with additional pipetting error. Note the absolute bias is on the log scale. Smaller on the log scale means closer to 0 on the original scale.

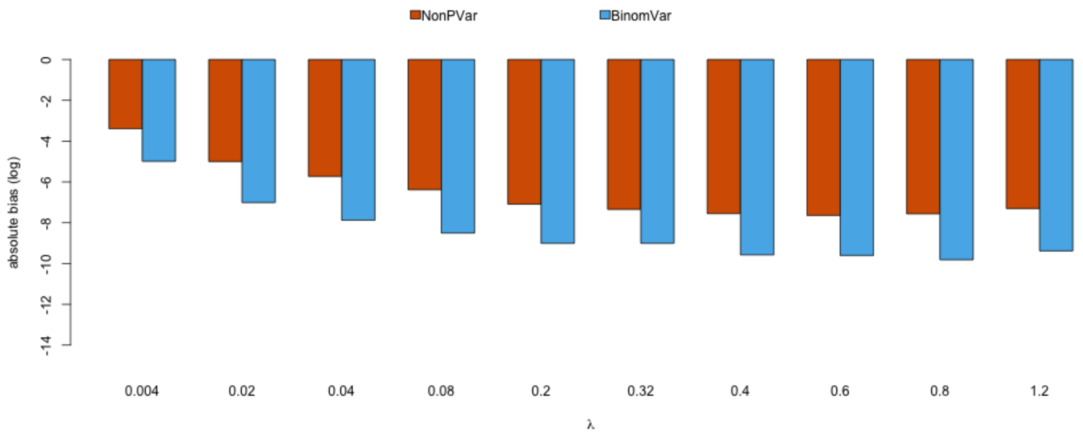

Figure S51: Absolute bias of the variance estimator of dsi in low DSI (=20%, 3 replicates) multiplex with additional partition loss. Note the absolute bias is on the log scale. Smaller on the log scale means closer to 0 on the original scale.

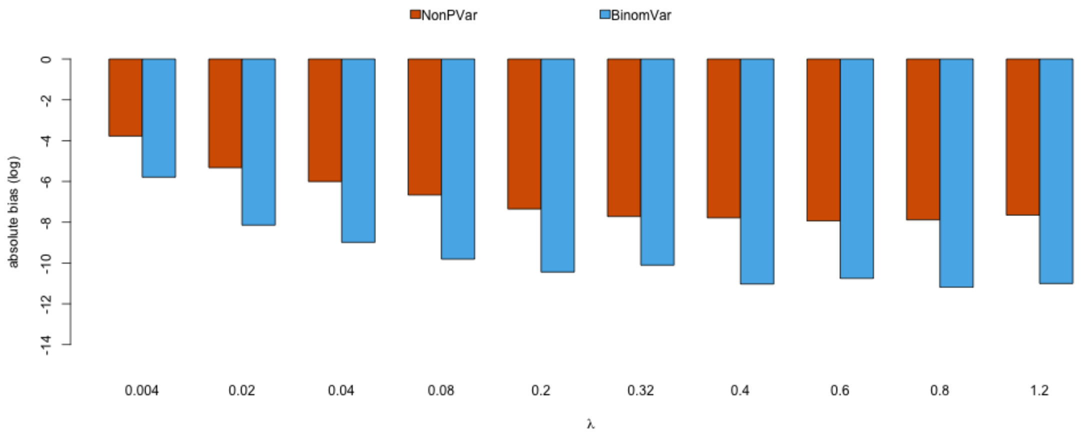

Figure S52: Absolute bias of the variance estimator of dsi in low DSI (=20%, 3 replicates) multiplex with additional partition size variation. Note the absolute bias is on the log scale. Smaller on the log scale means closer to 0 on the original scale.

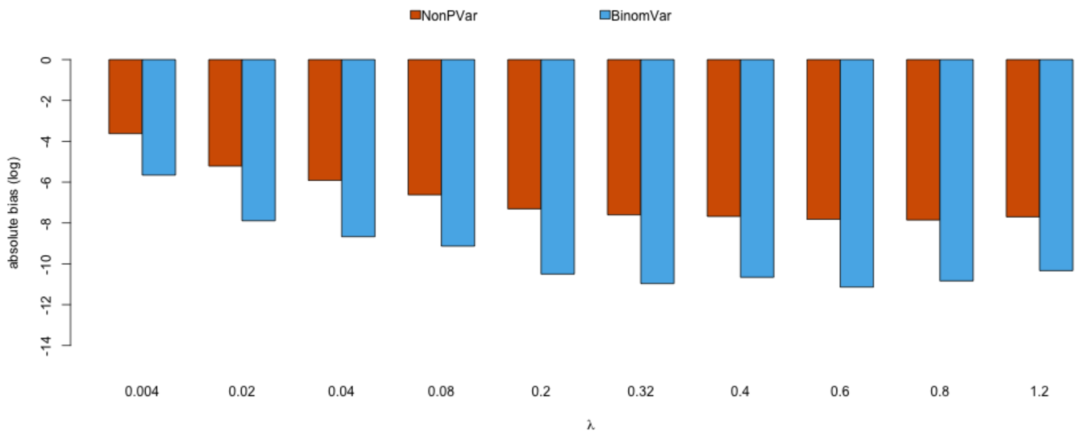

Figure S53: Absolute bias of the variance estimator of dsi in low DSI (=20%, 3 replicates) multiplex with additional misclassification. Note the absolute bias is on the log scale. Smaller on the log scale means closer to 0 on the original scale.

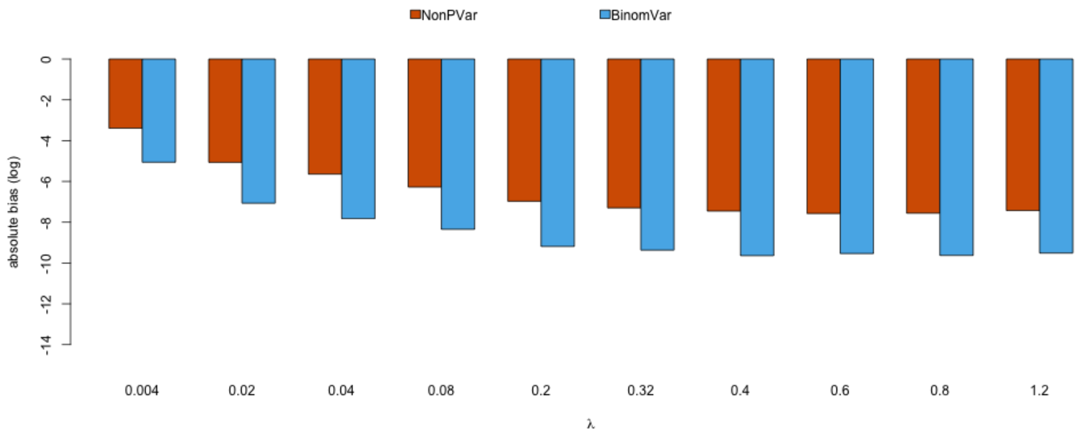

Figure S54: Absolute bias of the variance estimator of dsi in low DSI (=20%, 3 replicates) multiplex with all sources of variation. Note the absolute bias is on the log scale. Smaller on the log scale means closer to 0 on the original scale.

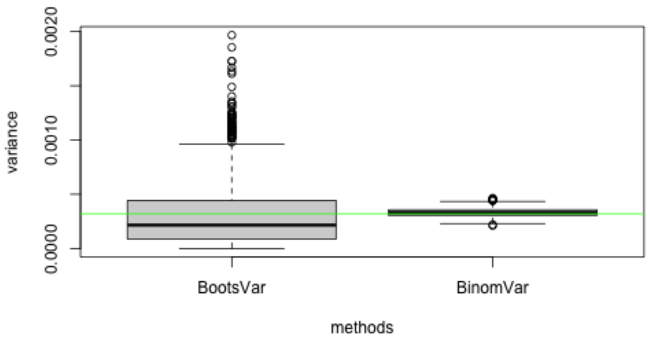

Figure S55: Variance estimates of simulation runs in low DSI (=20%, 3 replicates) multiplex with only sampling variation and random partitioning in low concentration setting of $\lambda_{AB} = 0.004$. The horizontal line is the estimated variance in the simulation (a good approximation to the true variance).

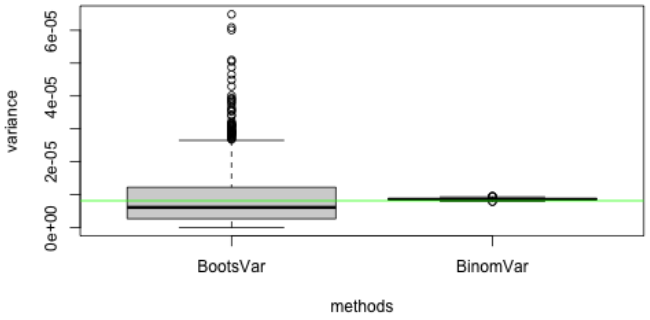

Figure S56: Variance estimates of simulation runs in low DSI (=20%, 3 replicates) multiplex with only sampling variation and random partitioning in medium concentration setting of $\lambda_{AB} = 0.2$. The horizontal line is the estimated variance in the simulation (a good approximation to the true variance).

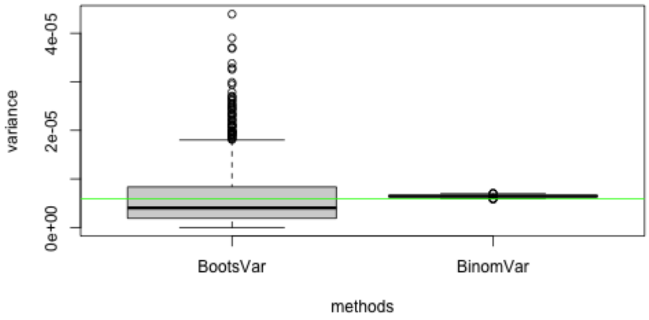

Figure S57: Variance estimates of simulation runs in low DSI (=20%, 3 replicates) multiplex with only sampling variation and random partitioning in high concentration setting of $\lambda_{AB} = 1.2$. The horizontal line is the estimated variance in the simulation (a good approximation to the true variance).

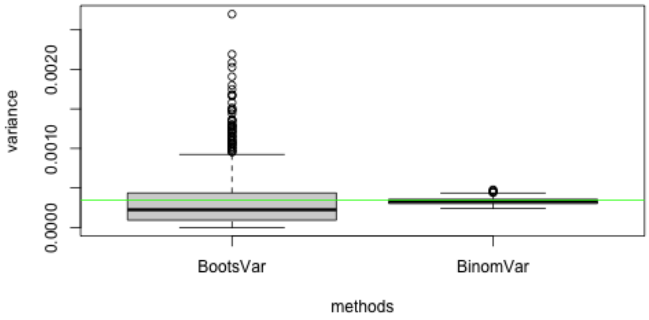

Figure S58: Variance estimates of simulation runs in low DSI (=20%, 3 replicates) multiplex with additional pipetting error in low concentration setting of $\lambda_{AB} = 0.004$. The horizontal line is the estimated variance in the simulation (a good approximation to the true variance).

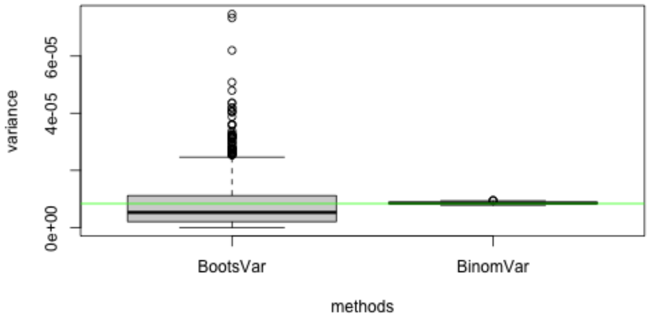

Figure S59: Variance estimates of simulation runs in low DSI (=20%, 3 replicates) multiplex with additional pipetting error in medium concentration setting of $\lambda_{AB} = 0.2$. The horizontal line is the estimated variance in the simulation (a good approximation to the true variance).

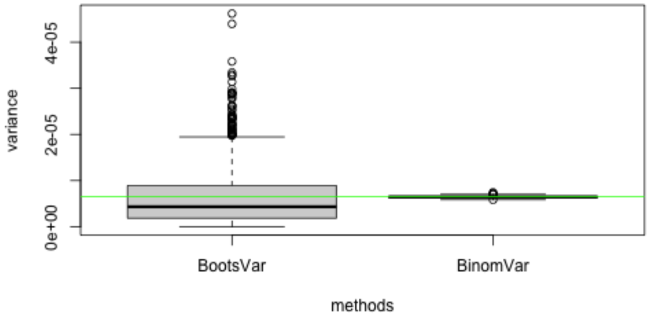

Figure S60: Variance estimates of simulation runs in low DSI (=20%, 3 replicates) multiplex with additional pipetting error in high concentration setting of $\lambda_{AB} = 1.2$. The horizontal line is the estimated variance in the simulation (a good approximation to the true variance).

#### 7.2 Medium DSI

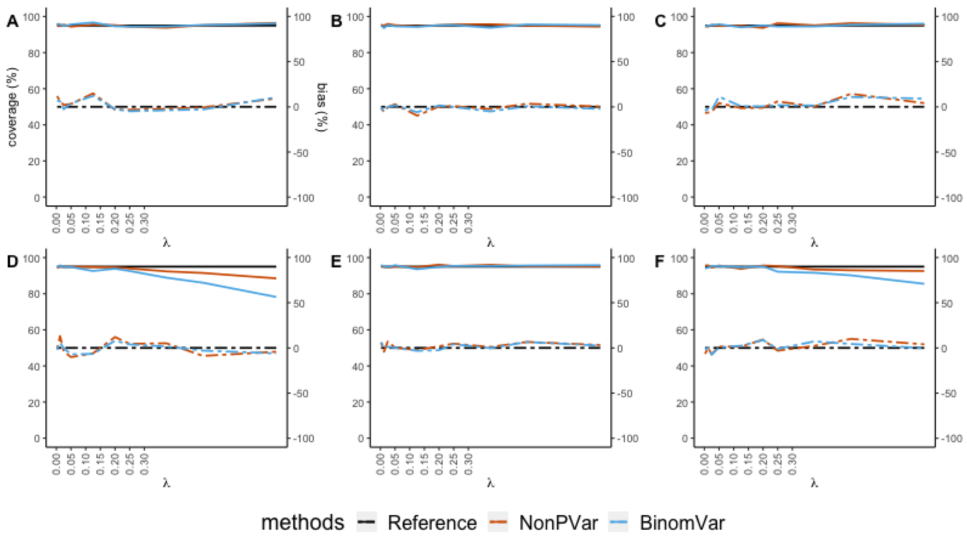

Figure S61: Empirical coverage of the 95% CIs (solid lines, left axis) and relative bias (dashed lines, right axis) for medium DSI (=50%, that is, 50% of the target molecules got sheared) in different scenarios. X-axis represents varying concentration of intact molecules from low to high. (A) only sampling variation and random partitioning with 3 replicates (B) 3% pipetting error (C) 20% partition loss (D) coefficient of variation of 10% in partition size (E) misclassification with 0.01% false positive rate and 5% false negative rate (F) all variation included. The reference (black solid line) for empirical coverage is set at 95%. The constructed CIs are supposed to cover the true values in 95% of the time. The closer other solid lines are to this reference, the better the CIs are. The reference (black dashed line) is set at 0%. The closer other dashed lines are to this reference, the lower the relative bias is.

##### 7.3 High DSI

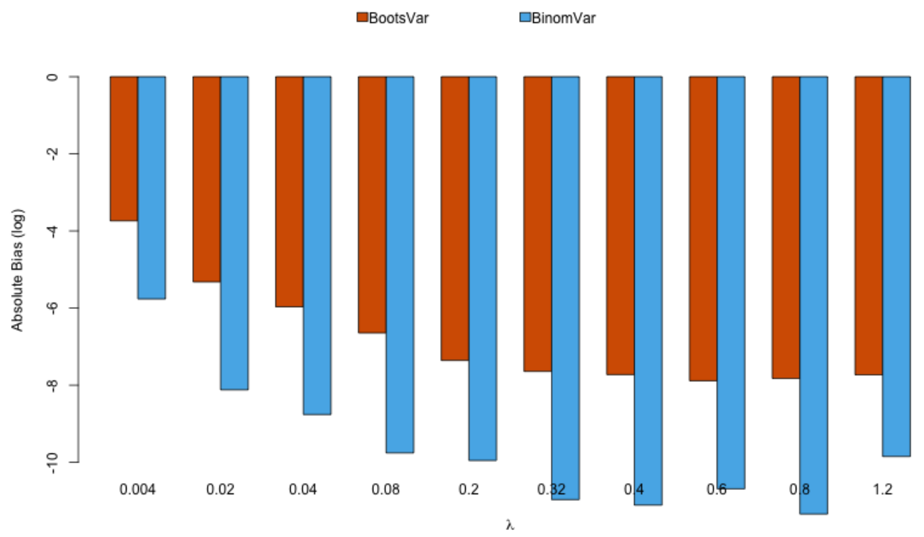

Figure S62: Empirical coverage of the 95% CIs (solid lines, left axis) and relative bias (dashed lines, right axis) for high DSI (=80%, that is, 80% of the target molecules got sheared) in different scenarios. X-axis represents varying concentration of intact molecules from low to high. (A) only sampling variation and random partitioning with 3 replicates (B) 3% pipetting error (C) 20% partition loss (D) coefficient of variation of 10% in partition size (E) misclassification with 0.01% false positive rate and 5% false negative rate (F) all variation included. The reference (black solid line) for empirical coverage is set at 95%. The constructed CIs are supposed to cover the true values in 95% of the time. The closer other solid lines are to this reference, the better the CIs are. The reference (black dashed line) is set at 0%. The closer other dashed lines are to this reference, the lower the relative bias is.

### 8 Real-life Data Analysis

CNV dataset is from Vynck et al. (2016) which consists of 10 samples with chromosomal abnormalities and 4 controls. For each sample and gene of interest, there are 2 or 3 technical replicates. The type of the data allows us to assess differences between Delta method, GLMM and the ones described in this paper.

Using the proposed framework in **Section CNV in Singleplex**, the copy numbers for each of the target loci was calculated by using the RPP30 locus as a reference.

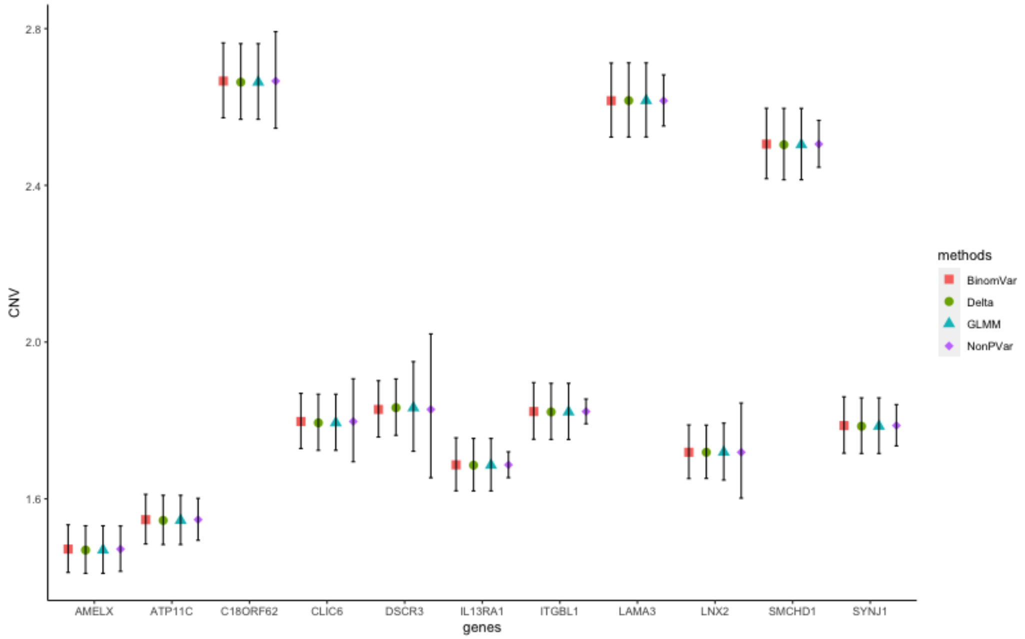

Figure S63: Copy numbers in sample 15 after normalization using the RPP30 locus (accounting for inter-replicate variability).

CNV is also constructed with delta method and GLMM models. Results in Fig.S63 show that the given estimates are quite close by all methods and the main difference is in the width of confidence interval. For some genes, the variance estimated by NonPVar is much smaller than the other 3 methods. That may indicate the variability is not reflected in those 3 replicates and more are required. Or it may indicate that the replicates are quite stable, thus NonPVar method gives a narrow confidence interval.

| **Methods** | **Runtime (s)** |
|---|---|
| **NonPVar** | 0.0065 |
| **BinomVar** | 0.0148 |
| **GLMM** | 13.4445 |
| **Delta Method** | 0.0066 |

Table S3: Computation time (4GB 1600 MHz DDR3) for 1 marker with 80% of partitions positive and 3 replicates. For BinomVar, the bootstrap iteration is set at 1000.

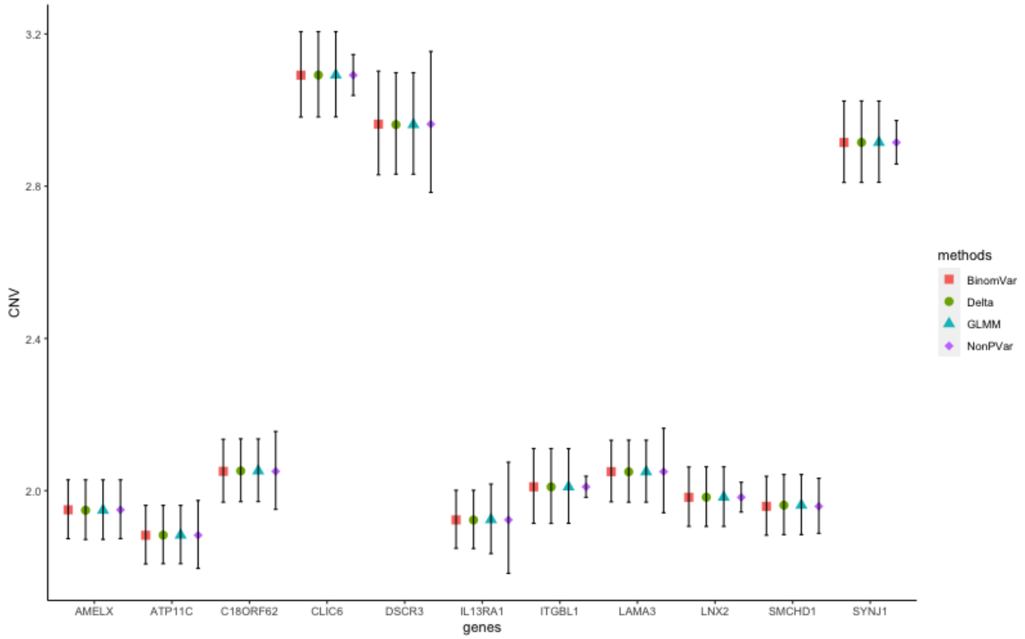

Figure S64: Copy numbers in sample 6 after normalization using the RPP30 locus (accounting for inter-replicate variability).
